# Phenotypic and genetic associations of quantitative magnetic susceptibility in UK Biobank brain imaging

**DOI:** 10.1101/2021.06.28.450248

**Authors:** Chaoyue Wang, Aurea B. Martins-Bach, Fidel Alfaro-Almagro, Gwenaëlle Douaud, Johannes C. Klein, Alberto Llera, Cristiana Fiscone, Richard Bowtell, Lloyd T. Elliott, Stephen M. Smith, Benjamin C. Tendler, Karla L. Miller

**Affiliations:** Wellcome Centre for Integrative Neuroimaging, FMRIB, Nuffield Department of Clinical Neurosciences, University of Oxford; Oxford Parkinson’s Disease Centre, University of Oxford, Oxford, UK; Donders Institute for Brain, Cognition and Behaviour, Centre for Cognitive Neuroimaging, Nijmegen, Netherlands; Sir Peter Mansfield Imaging Centre, School of Physics and Astronomy, University of Nottingham, Nottingham, United Kingdom; Department of Biomedical and Neuromotor Sciences, University of Bologna, Bologna, Italy; Department of Statistics and Actuarial Science, Simon Fraser University, Vancouver, BC, Canada

## Abstract

A key aim in epidemiological neuroscience is identification of markers to assess brain health and monitor therapeutic interventions. Quantitative susceptibility mapping (QSM) is an emerging MRI technique that measures tissue magnetic susceptibility and has been shown to detect pathological changes in tissue iron, myelin and calcification. We developed a QSM processing pipeline to estimate magnetic susceptibility of multiple brain structures in 35,885 subjects from the UK Biobank prospective epidemiological study. We identified phenotypic associations of magnetic susceptibility that include body iron, disease, diet, and alcohol consumption. Genome-wide associations related magnetic susceptibility to genetic variants with biological functions involving iron, calcium, myelin, and extracellular matrix. These patterns of associations include relationships that are unique to QSM, in particular being complementary to T2* measures. These new imaging phenotypes are being integrated into the core UK Biobank measures provided to researchers world-wide, creating potential to discover novel, non-invasive markers of brain health.

## Introduction

Magnetic resonance imaging (MRI) of the brain visualises anatomical structures on the scale of millimetres, but is sensitive to microscopic tissue features. This sensitivity confers the potential to detect the earliest stages of disease for therapeutic development and disease monitoring. A common challenge to this aim is a lack of risk factors that can be used to design cohorts targeting asymptomatic early disease; a powerful alternative is to prospectively image healthy individuals at large scale and track subsequent disease. The UK Biobank study is collecting brain imaging in 100,000 participants who are largely healthy when scanned^1^. Participants have been deeply phenotyped and genotyped, and consent to long-term access to their health records. UK Biobank has identified relationships between brain imaging markers and phenotypes including obesity^2^, vascular disease^3^ and ageing^4, 5^. It has also enabled major new insights into the genetic correlates of imaging phenotypes^6, 7^, identifying genes with known links to psychiatric illness^8, 9^, vascular disease^10, 11^ and neurodegeneration^12^.

UK Biobank is not yet fully exploiting the available brain imaging data, particularly the susceptibility-weighted MRI (swMRI) scan. swMRI signals are influenced by iron, myelin and calcium content due to the shifted magnetic susceptibility (χ) of these constituents relative to tissue water^13, 14^. The signal magnitude from swMRI has been analysed to provide estimates of signal decay time (T2*)^1, 15^, but the signal phase has not previously been analysed. Recently-developed algorithms for quantitative susceptibility mapping (QSM) transform swMRI phase data into quantitative estimates of χ^14, 16^. While derived from the same scan as T2*, QSM conveys distinct information. QSM estimates the mean χ within a voxel, reflecting bulk content of susceptibility-shifted sources like iron, whereas T2* reflecting the variance of χ-induced magnetic field fluctuations, reflecting compartmentalisation of these same sources. A consequence of this is that paramagnetic substances (e.g., iron) and diamagnetic substances (e.g., myelin) have the opposite effect on χ in QSM, but the same effect on T2*. QSM has been demonstrated to detect disease-relevant changes, such as iron accumulation in neurodegenerative disorders^17, 18^, and to provide an index of microstructural changes to tissue in normal ageing^19^. The UK Biobank brain thus creates unique opportunities to investigate QSM in previously unexplored territory, including as an early disease marker.

We developed a QSM pipeline for UK Biobank that was run on the current release of 35,885 participants, with repeat imaging in 1,447 participants. We conducted a comprehensive evaluation of established QSM algorithms and produced imaging-derived phenotypes (IDPs) of χ in a range of brain structures. We identified associations between these IDPs and non-imaging phenotypes, including diet, blood assays and health outcomes. We conducted the first genome-wide association studies (GWAS) using QSM-derived phenotypes, identifying relationships with genes with known relevance to iron, myelin and calcium, as well as less readily interpretable associations. Importantly, we found that QSM and T2* had distinct patterns of associations despite being derived from a single swMRI scan. QSM had higher heritability and two-timepoint agreement than T2* IDPs. Our QSM processing is now being incorporated into the core UK Biobank brain imaging processing pipeline^15^ to provide spatial χ maps and IDPs to researchers worldwide. These results demonstrate the richness of information in QSM data and the added value of QSM to the UK Biobank resource.

## Results

### Data analyses

We conducted an extensive evaluation of existing algorithms for each QSM processing step to establish an automated QSM pipeline. The final pipeline is illustrated in **Fig. 1a.** Details of evaluations are given in Supplementary §1.1. Briefly, individual channel phase images for each echo are combined using MCPC-3D-S^20^, unwrapped using a Laplacian-based algorithm^21^, and the two echoes are combined with weighted averaging.^22^. Brain-edge voxels with extremely large phase variance (primarily near sinuses) are detected and excluded. Background fields are removed using the V-SHARP algorithm^23^. Finally, χ maps are calculated using iLSQR^24^ and referenced to cerebrospinal fluid in the lateral ventricles.

**Figure 1.**
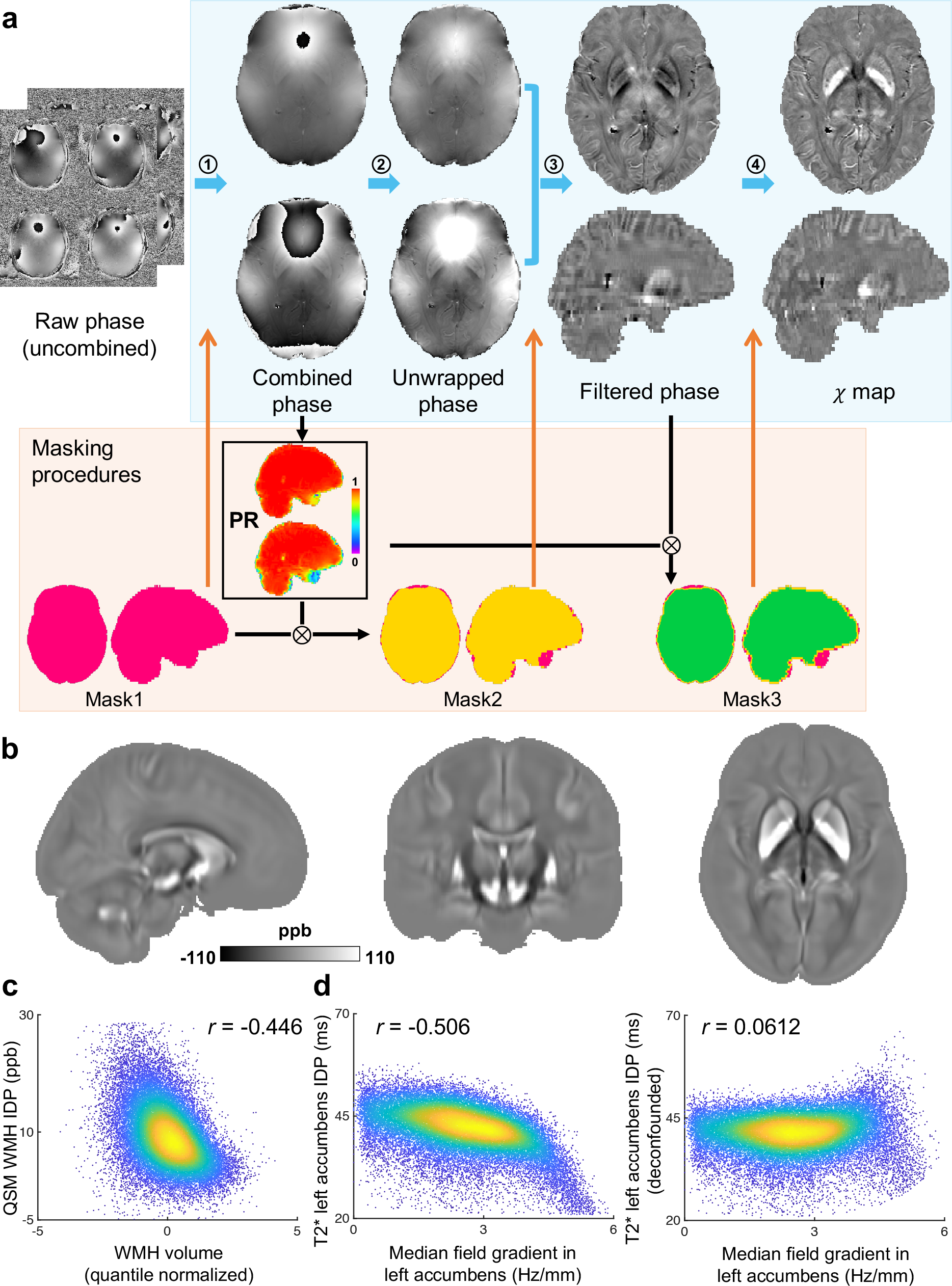
**(a)** QSM processing pipeline for UK Biobank swMRI data. Blue arrows indicate the main processing steps. Step 1: Channel combination using MCPC-3D-S. Step 2: Phase unwrapping using a Laplacian-based algorithm. Step 3: Background field removal using V-SHARP. Step 4: Dipole inversion using iLSQR. Black arrows indicate the brain mask evolution, and orange arrows indicate the brain mask applied at each step. Briefly, the brain mask provided by UK Biobank (Mask1, pink) was first used for the channel combination step. To exclude unreliable voxels in the vicinity of sinus cavities, the mask was subsequently refined using a ‘phase reliability’ map (PR, black box) (Mask2, yellow). After background field removal, the output mask from V-SHARP was further refined using the phase reliability map, with the resulting mask (Mask3, green) used for dipole inversion. Full details about the pipeline are provided in Methods. **(b)** QSM atlas generated by averaging χ maps (non-linearly registered to MNI space) from 35,273 subjects. **(c)** Association between QSM WMH IDP and WMH volume IDP (*r*=-0.446). **(d)** Example association between T2* left accumbens IDP and median field gradient measured in the left accumbens before (r=-0.506) and after (r=0.0612) deconfounding based on a physical model (details in Supplementary §2).

The pipeline was run on the 35,273 subjects with usable swMRI data, and each subject’s χ map was transformed to MNI standard space. A QSM population-average template (**Fig. 1b**) was produced by averaging all χ maps, and an “aging” template (**Supplementary Fig. S23b,d**) was calculated as the difference between the average χ maps for youngest (<52yo) and oldest (>75yo) subjects.

Median T2* IDPs in 14 major subcortical grey matter regions (accumbens, amygdala, caudate, hippocampus, pallidum, putamen and thalamus, both left and right) are already available in the current UK Biobank data release. We produced equivalent QSM-based IDPs, calculated as median χ values in these same 14 subcortical masks. We additionally extracted the median χ or T2* in the substantia nigra (left and right), bringing the total number of subcortical regions for both χ and T2* to 16. The UK Biobank also provides masks of white matter hyperintensities (WMH) derived from the T2-weighted structurals, which we used to derive the difference in χ or T2* between white matter hyperintensity (WMH) lesions and normal-appearing white matter^15^.

We observed strong correlations between WMH IDPs (both QSM and T2*) and WMH volume (**Fig. 1c**). Due to the relatively thick slices in the swMRI data, this correlation could reflect partial volume confounds. QSM and T2* WMH IDPs were thus additionally processed by regressing out the WMH volume, resulting in a total of 18 IDPs (16 subcortical and 2 WMH IDPs) for QSM or T2*.

We incorporated a new confound regressor based on a physical model^25^ that accounts for biases in T2* estimates introduced by macroscopic field gradients (Supplementary §2). As expected, this confound regressor correlated significantly with T2* IDPs (**Fig. 1d**) but not QSM (**Supplementary Fig. S16**), and as such was incorporated into phenotypic and genetic associations using T2*, but not QSM IDPs.

A reproducibility analysis using the two-timepoint data from 1,447 subjects demonstrated that subcortical QSM IDPs generally showed higher cross-scan correlation r compared with corresponding T2* IDPs (**Supplementary Fig. S10b**), particularly in the putamen, caudate, pallidum and substantia nigra (! > 0.8 for QSM and 0.6 < ! < 0.8 for T2*).

### Associations between IDPs and non-imaging phenotypes

We carried out univariate (pairwise) association analyses between 17,485 UK Biobank non-imaging phenotypes and QSM/T2* IDPs. For the remainder of this manuscript, we refer to all non-imaging phenotypes as “phenotypes”, to distinguish them from IDPs. These phenotypes have been grouped into 17 categories including early life factors (e.g., birth weight, maternal smoking), lifestyle (e.g., diet, alcohol consumption), physical/body measures (e.g., BMI, blood assays), cognitive test scores (e.g., numeric memory), health outcomes (e.g., clinical diagnosis – ICD10) and mental health variables (e.g., major depression).

The full set of 629,460 (17,485 phenotypes × 36 IDPs) correlations was corrected for multiple comparisons. We follow the convention for Manhattan plots and display results using -log10*P* (**Fig. 2**). Bonferroni correction for family-wise error (FWE) control at *Pcorrected*<0.05 was applied, corresponding to a -log10*Puncorrected* of 7.10. Additionally, a less conservative option for multiple comparison correction is false discovery rate (FDR), which for a 5% FDR resulted in -log10*Puncorrected*>3.13. In this manuscript, we primarily focus on associations passing the Bonferroni-corrected threshold, according to which we identified statistically-significant associations of 251 phenotypes with QSM IDPs, and 224 phenotypes with T2* IDPs. The total number of significant associations is much larger than this, as this count pools multiple time- point measurements of the same phenotype, and multiple IDPs associating with the same phenotype. The full list of significant phenotypic associations is provided in **Supplementary Table S1**.

**Figure 2.**
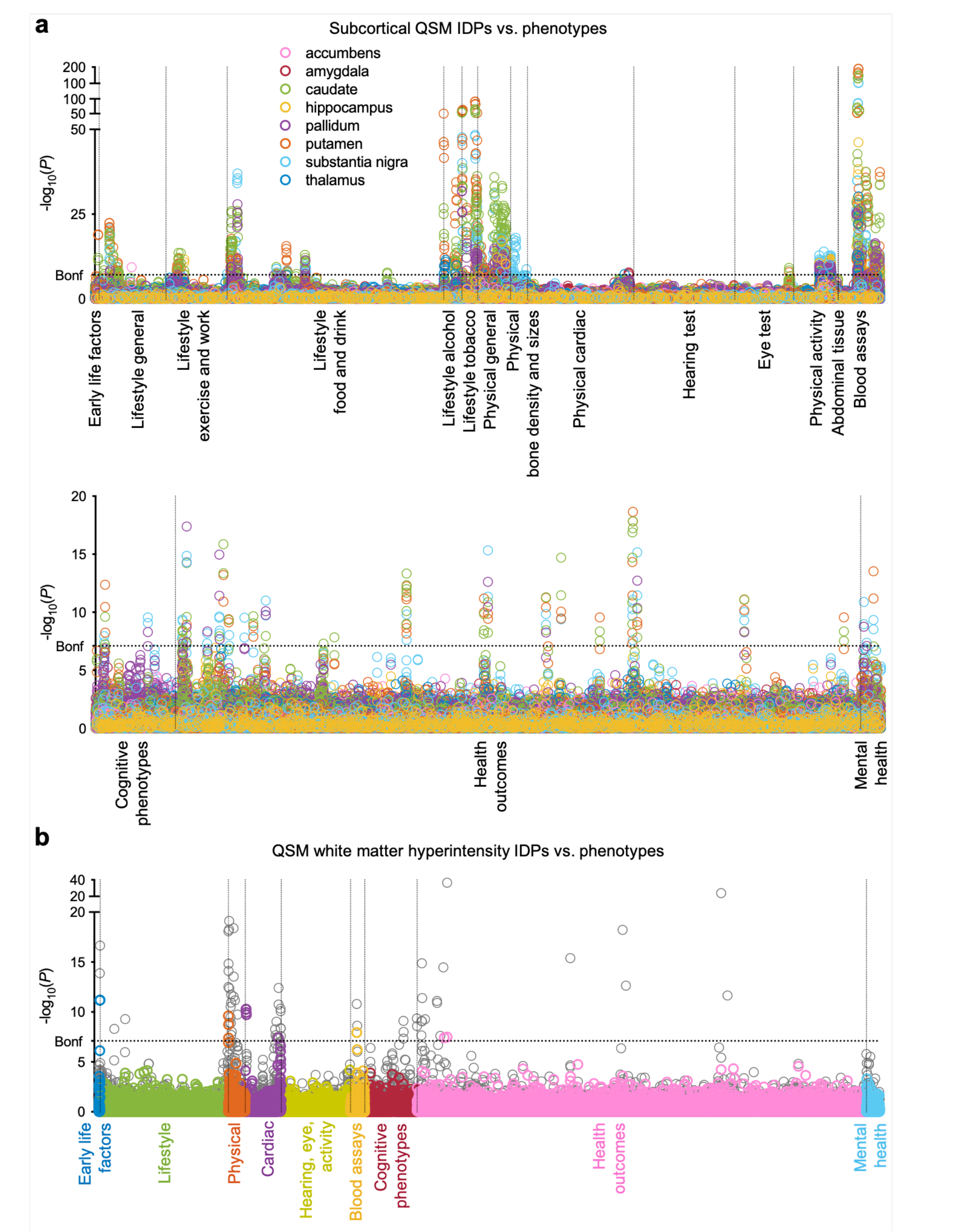
Visualization of univariate cross-subject association tests between 18 QSM IDPs and the 17,485 phenotypes in UK Biobank. Each circle represents a single IDP-phenotype association. The dashed horizontal line indicates the –log10*P* Bonferroni threshold of 7.10. All associations above this line are considered significant. Dashed vertical lines are used to distinguish between different phenotype categories **(a)** Manhattan plot showing associations between 16 subcortical QSM IDPs and phenotypes in 17 categories. **(b)** Manhattan plot showing associations between the QSM white matter hyperintensity (WMH) IDPs and all phenotypes (separated into 9 major categories). Shown behind (grey) are the associations without regressing out WMH volume.

**Table 1.** Summary of association results in 4 phenotype categories

| Phenotype category | Number of phenotypes | Brain regions (IDPs) | Phenotype/IDP associations used to derive voxel-wise maps in Fig. 4 |
| --- | --- | --- | --- |
| Blood assays | 33 | all 16 subcortical regions, WMH IDPs | *mean corpuscular haemoglobin* vs $\chi$ in right putamen ($r = 0.16$, $-\log_{10}P=190.37$) |
| Health outcomes | 44 | *MS*: thalamus, WMH IDPs;<br>*anaemia*: putamen, caudate, substantia nigra;<br>*diabetes*: pallidum, substantia nigra, caudate, putamen | *multiple sclerosis (self-reported)* vs $\chi$ in right thalamus ($r = -0.028$, $-\log_{10}P=6.85$);<br>*Anaemia (ICD10)* vs $\chi$ in left putamen ($r = -0.048$, $-\log_{10}P=18.65$);<br>*diabetes diagnosed by doctor* vs $\chi$ in right pallidum ($r = 0.047$, $-\log_{10}P=17.38$) |
| Food and drink | 22 | substantia nigra, pallidum, caudate, putamen, hippocampus | *tea intake* vs $\chi$ in right substantia nigra ($r = -0.069$, $-\log_{10}P=37.01$) |
| Alcohol consumption | 10 | putamen, caudate, thalamus | *frequency of consuming six or more units of alcohol* vs $\chi$ in thalamus ($r = -0.043$, $-\log_{10}P=9.94$) |

We compared the strength of QSM and T2* associations for each phenotype category (full results in Supplementary §3). Associations in some phenotype categories (e.g., *alcohol consumption*) are more specific to QSM IDPs (**Fig. 3a, b**) whereas other categories (e.g., *cardiac*) are more specific to T2* IDPs (**Fig. 3c, d**). However, the majority of phenotype categories show a mixed pattern of associations, including both common and distinct associations (e.g., *blood assays*) (**Fig. 3e**). This overall picture agrees with the expectation that QSM and T2* measures do not trivially recapitulate the same tissue properties, but together provide rich information from a single scan.

**Figure 3.**
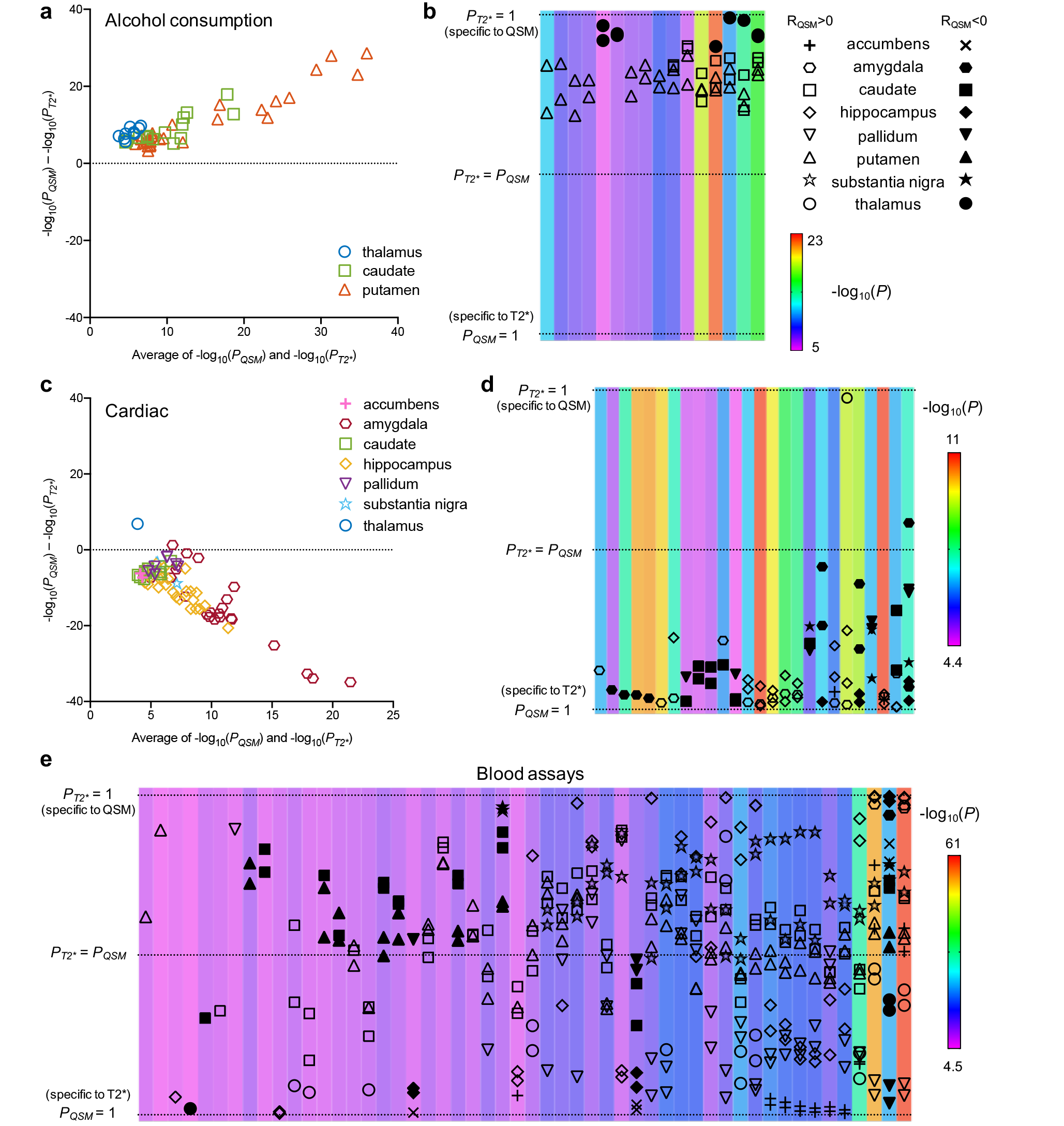
Example comparisons of phenotypic associations with QSM and T2* subcortical IDPs (ROI/phenotype pair shown if *PQSM* or *PT2\** passed the Bonferroni-corrected threshold). Here, we display results for **(a,b)** alcohol consumption, **(c,d)** cardiac and **(e)** blood assays categories. **(a,c)** Bland-Altman plot showing comparisons of -log10*P* values for QSM and T2* associations with (a) alcohol consumption and (c) cardiac categories. **(b,d,e)** Transformed Bland-Altman plot that aims to emphasise whether a given association is specific to QSM, T2*, or common to both. Each column represents one unique phenotype from the corresponding Bland-Altman plot, ordered from left to right by the number of associated regions. The vertical-axis is given by the angle of each point in a Bland-Altman plot with respect to the y=0 line. Hence, datapoints at the top (or bottom) of the plot represent an association that is highly specific to QSM (or T2*), and datapoints in the middle are phenotypes that associate with both QSM and T2* in a given brain region. The background colour of each column represents the average -log10*P* for significant associations with that phenotype. Unlike the Bland-Altman plot, this visualisation emphasises the modality specificity over the strength of correlation. For example, it is more apparent in (b) compared to (a) that thalamus-alcohol associations are highly specific to QSM. Here, the three categories reveal more QSM-specific (a,c), T2*-specific (b,d) and mixed (e) association patterns.

Having identified phenotypes that associate with QSM IDPs, we conducted voxel-wise regressions with these same phenotypes into χ maps to investigate the spatial regions driving these associations. **Figure 4** shows voxel-wise associations of χ with 6 representative phenotypes. Voxel-wise association maps with lead associations in each phenotype category, are provided in Supplementary §4. In general, voxel-wise association maps are highly symmetric, including extended homogeneous regions, more focal associations with sub- regions, and associations with brain areas not included in the ROIs used to generate QSM IDPs.

**Figure 4.**
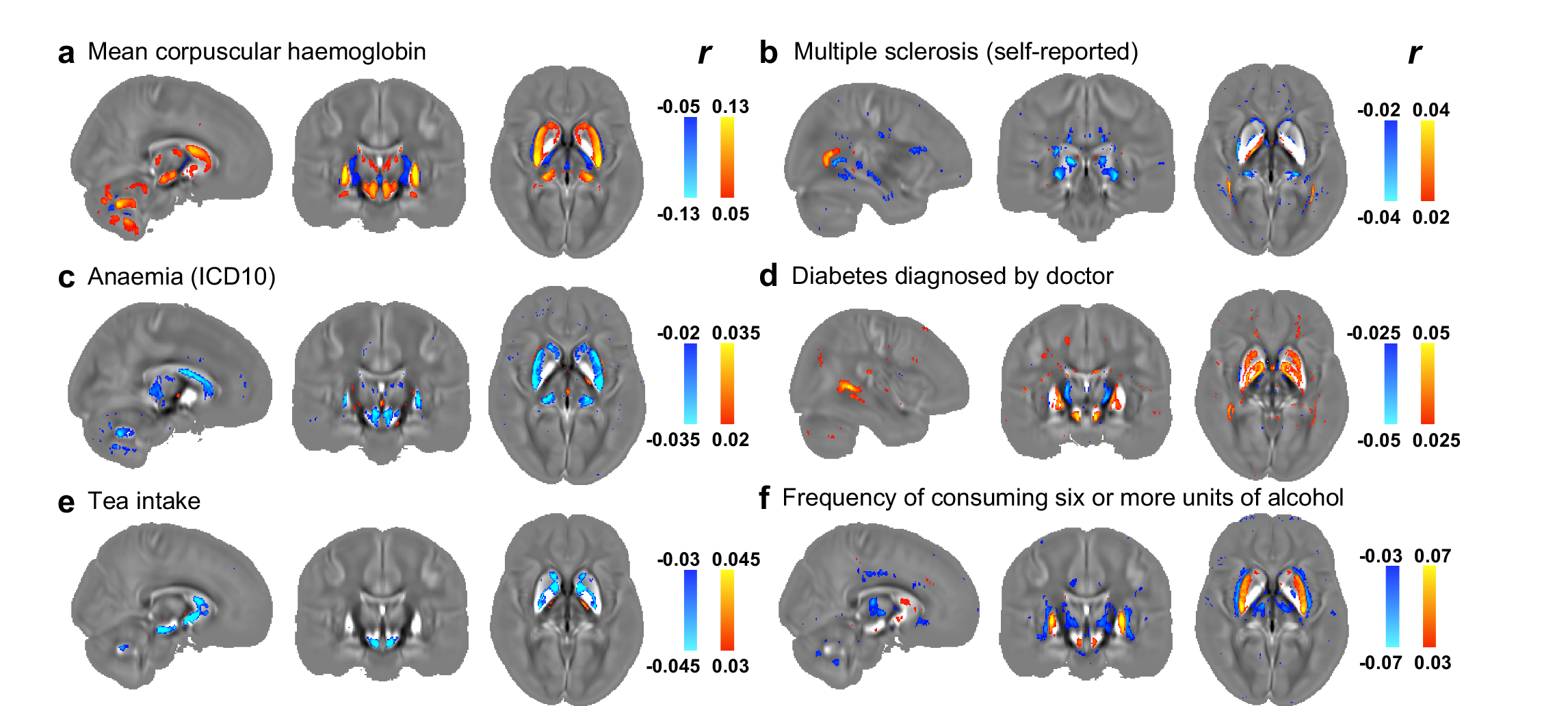
Voxel-wise association maps of 6 example phenotypes with χ maps aligned in MNI space. Pearson correlation r is shown as color overlay (red-yellow for positive r and blue for negative r) on the population-average χ map. **(a)** *mean corpuscular haemoglobin* identifies all subcortical regions captured by our IDPs, as well as the red nucleus and cerebellar regions. Particularly, the putamen, caudate, substantia nigra and red nucleus exhibit homogeneous correlations across the entire region; **(b)** *multiple sclerosis (self-reported)* identifies sub-regions of thalamus (including the pulvinar nucleus and lateral geniculate nucleus), as well as focal white matter regions such as the optic radiation; **(c)** *Anaemia (ICD10)* identifies putamen, caudate, red nucleus and cerebellar regions, as well as sub-regions of substantia nigra and thalamus; **(d)** *diabetes diagnosed by doctor* identifies sub-regions of caudate, putamen, pallidum and substantia nigra, in addition to white matter regions including the splenium of the corpus callosum and optic radiations; **(e)** *tea intake* identifies sub-regions of the caudate, pallidum and substantia nigra; **(f)** *frequency of consuming six or more units of alcohol* identifies putamen and sub-regions of thalamus, caudate and substantia nigra.

We now describe associations in specific phenotype categories in more detail, focusing on associations that recapitulate previous studies or are more specific to QSM IDPs. An overview of these categories is given in **Table 1**.

### Phenotypic associations with QSM IDPs in four categories

#### Blood assays

Phenotypic associations with blood assays include haemoglobin, cell counts, cell morphology and blood constituents. The strongest of these are haemoglobin-related phenotypes, which show strong, positive correlations with QSM IDPs (**Fig. 2a** and **Supplementary Table S1**). The voxel-wise map of association with *mean corpuscular haemoglobin* (**Fig. 4a**) (the strongest phenotypic correlation) reveals spatially contiguous positive associations in all subcortical regions captured by our IDPs, as well as the red nucleus and cerebellar regions. Here, the putamen, caudate, substantia nigra and red nucleus exhibit homogeneous correlations across the entire region, while voxels in the pallidum, hippocampus and thalamus localise to specific sub-regions. Haemoglobin-related blood measures are used clinically as a marker for a subject’s iron level^26^, and the positive sign of associations with haemoglobin measures is consistent with both QSM’s established relationship with iron, and the positive correlation with iron concentration from post-mortem studies^27^. Associations with QSM WMH IDPs are distinct to subcortical regions, exhibiting specificity to red blood cells and haematocrit.

#### Health outcomes

QSM IDPs are associated with multiple sclerosis (MS), anaemia and diabetes. Multiple sclerosis is significantly associated with QSM WMH IDPs, which is in line with previous studies that reported altered χ in MS lesions^28^. Previous literature reported decreased χ in the thalamus (particularly pulvinar nucleus) for MS patients compared to healthy volunteers^29^. Associations between QSM right thalamus IDP and *multiple sclerosis (self-reported)* (! = −0.028, -log10*P*=6.85) is significant at the FDR-corrected threshold (-log10*P*=3.13), but is just below the Bonferroni-corrected threshold (-log10*P*=7.1). The voxel-wise association map with *multiple sclerosis (self-reported)* (**Fig. 4b**) reveals spatially contiguous negative associations in sub-regions of thalamus (including the pulvinar nucleus and lateral geniculate nucleus), as well as focal white matter regions such as the optic radiation. This suggests that the sub- threshold association at IDP level may be due to the use of ROIs covering the entire thalamus that dilute significance of results that are specific to sub-regions. Interestingly, previous literature has reported structural damage of the thalamic lateral geniculate nucleus in MS patients^30^, reflecting potential damage of the visual pathway in MS. Associations with self-reported and diagnosed anaemia are consistent with a reduction in tissue iron^27^, finding negative correlations with χ and positive correlations with T2*. The voxel-wise association map with *anaemia (ICD10)* (**Fig. 4c**) reveals spatially contiguous negative associations in the putamen, caudate, red nucleus and cerebellar regions, as well as sub-regions of substantia nigra and thalamus. QSM associations with diabetes include formal diagnosis, self-report, and relevant medication (insulin and metformin). The voxel-wise association map with *diabetes diagnosed by doctor* (**Fig. 4d**) reveals spatially contiguous positive associations in the caudate, putamen, pallidum and substantia nigra regions, in addition to white matter including the splenium of the corpus callosum and optic radiations. Body iron overload in diabetes has been frequently reported^31^, with a recent brain imaging study finding increased χ in the caudate, putamen and pallidum in type 2 diabetes^32^, in agreement with our results. Finally, QSM WMH IDPs correlated with hypertension and measures of blood pressure; vascular risk factors (including hypertension) have been reported to have an effect on MS pathology^33^ which may result in changes of χ in WMH lesions.

#### Food and drink

QSM IDPs are associated with food and drink intake include tea, coffee, meat and carbohydrate consumptions, as well as dietary supplements (**Supplementary Table S1).** The strongest associations relate to tea consumption, which correlates negatively with χ. Although no direct link between tea intake and χ measures have been described previously, polyphenols in both green and black tea have been reported as brain-permeable, natural iron chelators that have demonstrated neuroprotective effects^34, 35^. The voxel-wise association map with *tea intake* (**Fig. 4e**) reveals spatially contiguous negative associations in sub-regions of the caudate, pallidum and substantia nigra. This spatial pattern of correlations is in line with a previous rodent study in which black tea extract reduced oxidative stress levels in the substantia nigra and striatum^34^.

### Alcohol consumption

Alcohol consumption correlated more strongly with QSM IDPs than T2* IDPs in all cases (**Fig. 3a, b**). Voxel-wise association map with *frequency of consuming six or more units of alcohol* (**Fig. 4f**) reveals spatially contiguous positive associations in the putamen and sub-regions of substantia nigra and caudate, but also negative associations in sub-regions of the thalamus. These results recapitulate a previous study finding higher χ in the putamen, caudate and substantia nigra in subjects with alcohol use disorder^36^ which has been linked to abnormal body iron accumulation^37, 38^. χ in the thalamus correlated with phenotypes relating to the quantity of alcohol consumption, in some cases (e.g., *frequency of consuming six or more units of alcohol*) having no significant correlation with T2*. The thalamus is involved in the frontocerebellar circuit and Papez circuit, which are particularly affected by alcohol consumption^39, 40^. Although no previous studies have linked χ in the thalamus with alcohol use disorder, neuroimaging studies have reported reductions in thalamic volume and connectivity in alcohol use disorder patients^39, 40^.

### Associations between IDPs and genetic variants

#### Heritability of QSM and T2* IDPs

Following previous studies^6, 7^, we use linkage score regression^41^ to estimates narrow sense heritability^42^ (*h*^2^) as the fraction of IDP variance that is explained by a linear combination of genetic variants. *h*^2^ ranges from 0 (independent of genotype) to 1 (entirely determined by genotype). Subcortical QSM and T2* IDPs are highly heritable, being more than one standard error >0 (**Fig. 5a**). In all but one IDP, QSM has higher heritability than T2*. χ in the putamen and substantia nigra showed the highest heritability (0.323-0.342), while T2* in the amygdala and accumbens showed two of the lowest heritability estimates (0.0357-0.0684). Heritability of all brain IDPs in UK Biobank were previously reported in the range of 0.000-0.405^7^. The heritability of χ in right putamen (*h*^2^=0.342) is >98% among UK Biobank brain IDPs, and roughly half the heritability of human height^42^.

**Figure 5.**
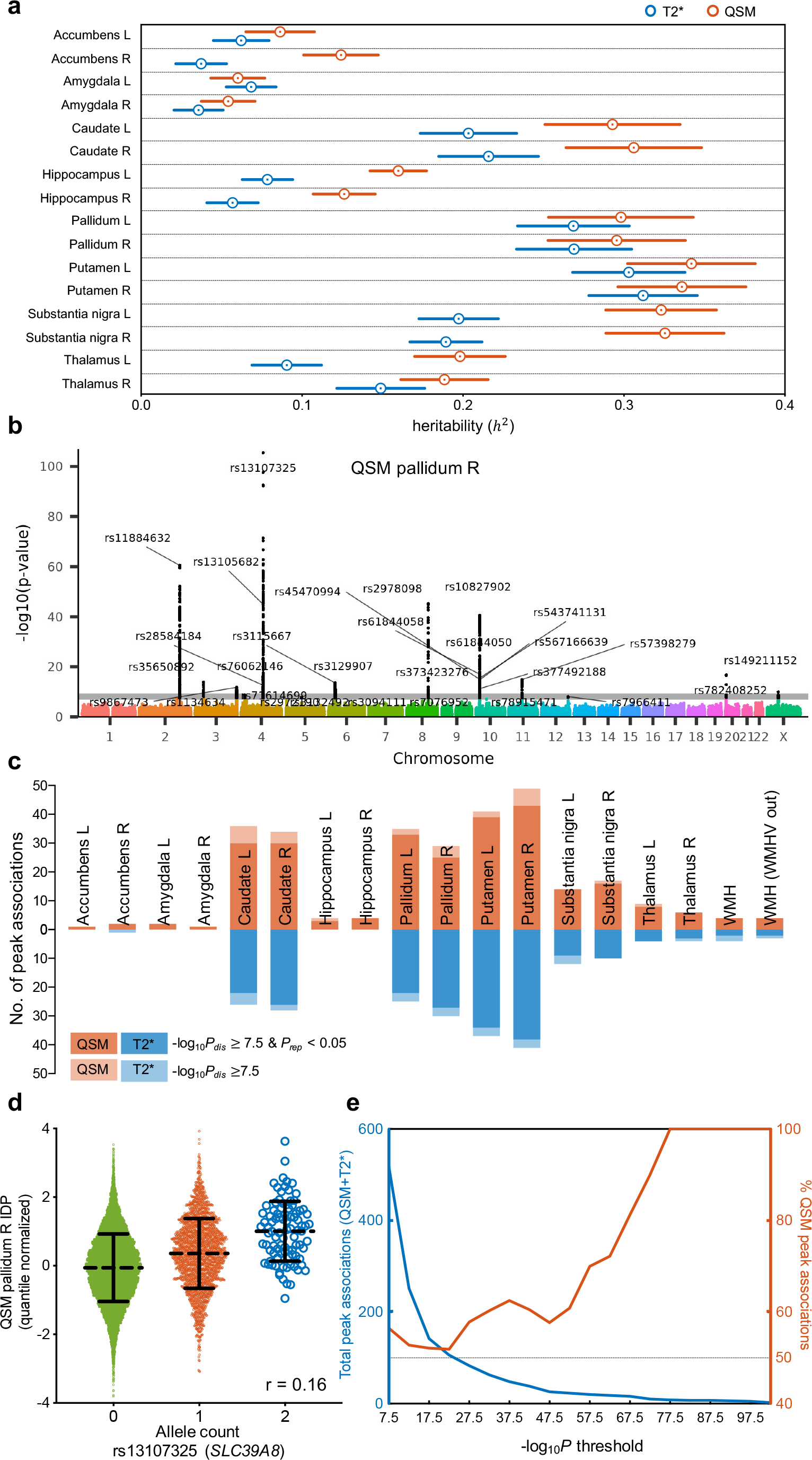
**(a)** Heritability estimates (*h*^2^) for subcortical QSM and T2* IDPs grouped according to regions. Circle indicates heritability estimate and error bar indicates standard error. **(b)** Example Manhattan plot relates to the GWAS for QSM right pallidum IDP. The lower grey horizontal line indicates the -log10*P* threshold of 7.5 and the upper line the Bonferroni threshold of 9.06. **(c)** Stacked bar chart showing comparisons of number of peak associations identified in GWASs (passing the -log10*P* threshold of 7.5) for QSM vs T2* IDPs. **(d)** Scatterplot showing the relationship between QSM right pallidum IDP vs allele count of rs13107325 (the strongest genetic association across all GWASs). **(e)** Distribution of -log10*P* values of all peak associations identified in GWASs (blue line, left y axis). Right y axis (orange line) is showing percentage of peak associations identified with QSM IDPs.

#### Genome-wide associations studies of IDPs

We carried out a GWAS for each QSM and T2* IDP following a previously-described approach^6, 7^ using the second release of over 90 million imputed genetic variants. Subjects were divided into discovery (n=19,720) and replication (n=9,859) cohorts. The standard single-phenotype GWAS threshold (-log10*P*=7.5) and also a more stringent threshold after additional Bonferroni correction to account for the number of GWASs (18 × 2) carried out (resulting in a Bonferroni threshold of -log10*P*=9.06) were used. We report the genetic variant with the strongest “peak” association in each region of linkage disequilibrium (LD, see Methods). **Figure 5b** displays a Manhattan plot for QSM right pallidum IDP. In total, QSM IDPs identified 292 peak associations (265 replicated), T2* IDPs identified 225 peak associations (199 replicated). **Figure 5c** provides a summary of peak associations from the set of GWASs. The strongest genetic association across all GWASs was found between QSM right pallium IDP and variant rs13107325, which is shown in **Fig.5d**. **Figure 5e** provides a summary of the distribution of -log10*P* values of all peak associations identified in GWASs. Supplementary §5 includes Manhattan plots for all GWASs and **Supplementary Tables S2-3** provides the full list of peak associations.

**Table 2.**
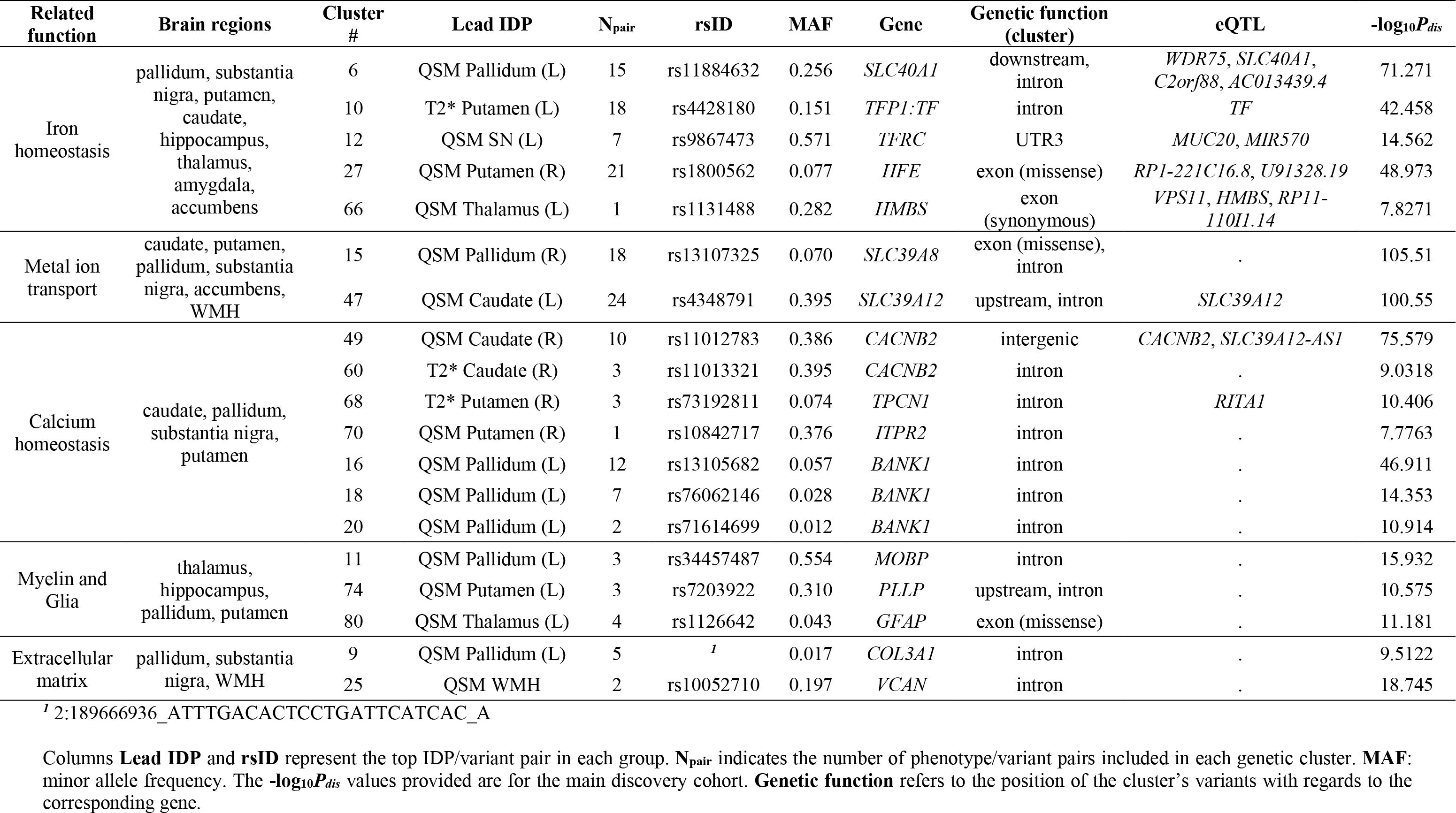
Information for the 19 example genetic clusters

We used the Peaks software^7^ (https://github.com/wnfldchen/peaks) to automatically generate clusters of peak associations between genetic variants and IDPs. A cluster is defined using the discovery cohort as a set of IDP-variant pairs for which all genetic variants are within a 0.25- cM distance of the top variant within the cluster. We classify a cluster as replicating if at least one of its IDP-variant pairs had nominal significance (*P*<0.05) in the replication cohort. We used FUMA^43^ to map the genetic variants of each cluster to related genes.

We identified 89 distinct clusters, 80 of which replicated. Among the replicated clusters, 54 had associations with both QSM and T2* IDPs, 22 were unique to QSM IDPs, and 4 were unique to T2* IDPs. All clusters common to QSM and T2* IDPs replicated. Note that a cluster can include just a single genetic variant, which was the case for 11 replicated clusters. **Table 2** provides a summary of 19 example clusters (the full list of clusters is given in **Supplementary Table S4)**. Most replicating clusters are associated with genes, including 10 clusters with variants in exons (6 missense). Many clusters are associated with genes involved in functions with known relevance to tissue χ, including myelination, iron and calcium. Other clusters were associated with genes whose function does not have an expected relationship to tissue χ, including transcription factors, extracellular matrix, and intracellular trafficking. Below, we describe select examples in detail.

#### Iron transport and homeostasis

Multiple clusters are related to genes implicated in iron transport and homeostasis. **Cluster 6** comprises eight genetic variants related to the ferroportin gene (*SLC40A1*). Voxel-wise association map with rs11884632 (*SLC40A1*, **Fig. 6a**) includes pallidum, sub-regions of substantia nigra and thalamus, red nucleus and cerebellar nuclei. Ferroportin exports iron from cells, and mutations in *SLC40A1* lead to hemochromatosis^44^. Three functionally-related clusters identified associations with the transferrin gene (*TF*, **cluster 10**), the transferrin receptor gene (*TFRC*, **cluster 12**), and the homeostatic iron regulator gene (*HFE*, **cluster 27**, including a missense variant). Voxel-wise association map with rs1800562 (*HFE*, **Fig. 6b**) includes putamen, red nucleus, cerebellar regions, sub-regions of caudate, substantia nigra, and thalamus. Voxel-wise association map with rs4428180 (*TF*) shows similar pattern of association to rs1800562 (*HFE*), and rs9867473 (*TFRC*) shows similar pattern of association to rs11884632 (*SLC40A1*) (**Supplementary Fig. S26**). The TF protein delivers iron to proliferating cells via TFRC, an interaction that is modulated by the HFE protein to regulate iron absorption. Mutations in *HFE* lead to hereditary hemochromatosis, while mutations in *TF* lead to hereditary atransferrinemia^44^. The variants we identified have been previously associated with transferrin levels^45^, iron biomarkers^46^, and Alzheimer’s disease^47^. Interestingly, associations with variants related to *SLC40A1* (iron export) and *HFE* (iron absorption) had opposite signs in corresponding regions, in line with their biological functions (**Fig. 6a,b**). **Cluster 66** comprises a single association of QSM in left thalamus with a potentially deleterious exonic variant (synonymous, CADD: 17.8) in the *HMBS* gene. The voxel-wise association map with this variant (rs1131488, *HMBS*) identifies sub-regions of thalamus and dispersed white matter (**Fig. 6c**). *HMBS* encodes an enzyme from the heme biosynthetic pathway. *HMBS* mutations are associated with acute intermittent porphyria^48^ and leukoencephalopathy, which exhibit MRI anomalies in thalamus and cerebral white matter^49^.

**Figure 6.**
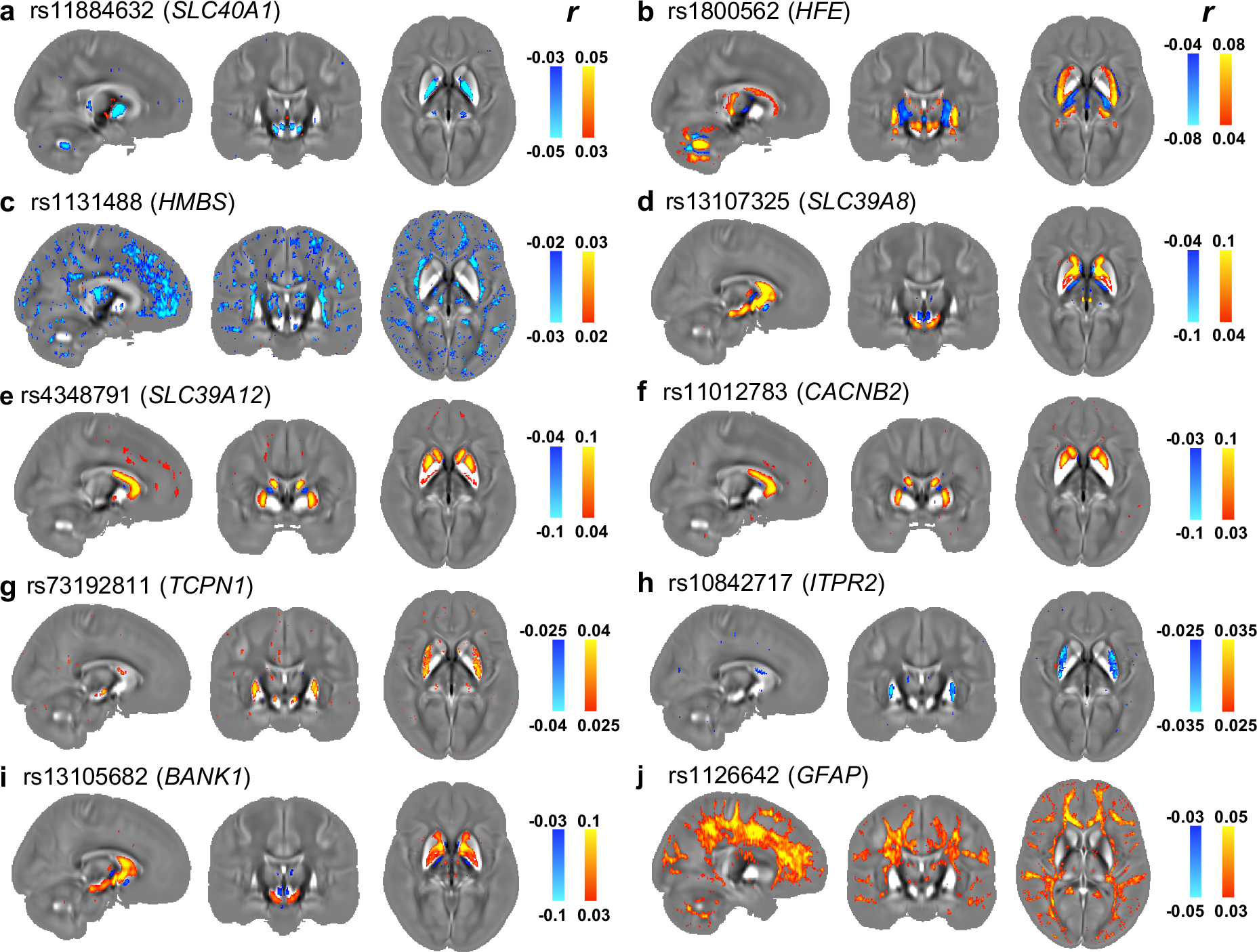
Voxel-wise association maps of top genetic variants of 10 genetic clusters with χ maps aligned in MNI space. Pearson correlation r is shown as color overlay (red-yellow for positive r and blue for negative r) on the population-average χ map. **(a)** rs11884632 (*SLC40A1*) identifies pallidum, sub-regions of substantia nigra and thalamus, red nucleus and cerebellar nuclei; **(b)** rs1800562 (*HFE*) identifies putamen, red nucleus, cerebellar regions, sub-regions of caudate, substantia nigra, and thalamus; **(c)** rs1131488 (*HMBS*) identifies sub-regions of thalamus and dispersed white matter; **(d)** rs13107325 (*SLC39A8*) identifies caudate, substantia nigra and sub- regions of pallidum; **(e)** rs4348791 (*SLC39A12*) identifies caudate and sub-regions of putamen and pallidum; **(f)** rs11012783 (*CACNB2*) identifies caudate and sub-regions of putamen; **(g)** rs73192811 (*TPCN1*) identifies putamen and sub-regions of substantia nigra; **(h)** rs10842717 (*ITPR2*) identifies putamen; **(i)** rs13105682 (*BANK1*) identifies caudate, substantia nigra and sub-regions of pallidum; **(j)** rs1126642 (*GFAP*) identifies sub- regions of thalamus and widespread white matter regions.

#### Metal ion transporters

There are also clusters associated with genes from the SLC39 family of solute-carriers, which transport divalent metal cations such as Zn^2+^ and Fe^2+^. **Cluster 15** comprises four genetic variants associated with QSM and T2* IDPs in multiple subcortical structures and WMHs. The top variant (rs13107325) is a missense variant of *SLC39A8* (*ZIP8*). Voxel-wise association map with this variant (**Fig. 6d**) identifies caudate, substantia nigra and sub-regions of pallidum. *SLC39A8* encodes a transmembrane transporter protein for zinc and iron. This genetic variant has been linked to blood pressure^50^, diabetes^51^, Parkinson’s disease^52, 53^, schizophrenia^52, 53^, alcohol consumption^54^, haemoglobin and haematocrit^55^, and brain morphology^6, 56, 57^. **Cluster 47** comprises eleven genetic variants related to *SLC39A12* (*ZIP12*). The voxel-wise association map for the top variant (rs4348791, **Fig. 6e**) identifies caudate and sub-regions of putamen and pallidum. *SLC39A12* encodes a zinc/iron transmembrane transporter that is highly expressed in the brain; low expression of *SLC39A12* leads to impaired neural development^58^.

#### Calcium homeostasis

Seven clusters are related to calcium channels and regulation. **Clusters 49** and **60** contain variants related to *CACNB2*. Voxel-wise association map with rs11012783 (*CACNB2*) (**Fig. 6f**) identifies caudate and sub-regions of putamen. *CACNB2* encodes a subunit of voltage-gated calcium channels that regulate calcium influx from the extracellular space^59^. Variants in *CACNB2* have been associated with autism, bipolar disorder, depression and schizophrenia^60–62^. **Cluster 68** includes two intronic variants in *TPCN1* and **cluster 70** includes one genetic variant in an intron of *ITPR2*. Voxel-wise maps exhibit a similar spatial pattern in the putamen for rs73192811 (*TCPN1*, **Fig. 6g**) and rs10842717 (*ITPR2*, **Fig. 6h**), but with opposite signs. Both *TCPN1* and *ITP2* encode calcium channels that control the release of calcium from intracellular spaces. **Clusters 16**, **clusters 18** and **clusters 20** are related to *BANK1*. Voxel-wise association map with rs13105682 (*BANK1*) (**Fig. 6i**) reveals spatially contiguous associations in the caudate, substantia nigra and sub-regions of pallidum. *BANK1* encodes a protein that regulates calcium mobilization from intracellular stores that is primarily related to the immune system, but is also expressed in the brain. Variants in *BANK1* have been related to working memory task-related brain activation^63^.

#### Glia and myelin

We observed associations with genetic variants in two genes that encode structural constituents of myelin sheaths. **Cluster 74** includes genetic variants related to *PLLP*, which encodes the myelin protein plasmolipin, and **cluster 11** includes genetic variants located in an intron of *MOBP*, which encodes the myelin-associated oligodendrocyte basic protein. **Cluster 80** includes a missense variant of *GFAP*, which encodes an intermediate filament protein that is highly specific for cells of astroglial lineage. Voxel-wise association map with rs1126642 (*GFAP*) (**Fig. 6j**) reveals spatially contiguous associations in sub-regions of thalamus and widespread white matter regions. Mutations in *GFAP* lead to Alexander disease, a genetic disorder characterized by fibrinoid degeneration of astrocytes, demyelination and white matter anomalies^64^. *GFAP* variations have also been associated with white matter microstructure phenotypes and Alzheimer’s disease^57, 65, 66^. While astrocytes are present in both grey and white matter, *GFAP* expression is higher in white matter astrocytes than in grey matter astrocytes^67^.

#### Extracellular matrix

Many genetic associations do not have an obvious, direct link to magnetic susceptibility contrast. For example, **cluster 25** includes a single genetic variant (rs10052710) located in an intron of *VCAN* that is associated with QSM in WMHs (both with and without regressing out WMH volume). *VCAN* encodes versican, a major component of the extracellular matrix in multiple tissues including the brain, which is highly expressed in the brain during development^68^. In addition to its structural role, versican can also interact with inflammation and the immune response^69^, and its expression is altered in multiple sclerosis lesions^70, 71^. This and other variants in *VCAN* have been previously associated with multiple brain phenotypes, in particular to white matter microstructure metrics derived from diffusion MRI^6, 57, 72^.

## Discussion

### QSM in population imaging

Unlike the other brain imaging modalities in UK Biobank, which have been (or are being) collected or collated previously in thousands of subjects^73–76^, QSM has previously been limited to smaller-scale studies. This UK Biobank QSM resource is approximately 2 orders of magnitude greater than the largest existing QSM dataset^77^. The number of subjects, coupled with the breadth of linked data, including genetics, extensive phenotyping and health outcomes, is expected to open up new avenues of investigation for QSM. At the time of scanning, most UK Biobank subjects are largely healthy, with the cohort age range designed to reflect a broad range of health outcomes in the coming decades. Hence, this cohort is particularly appropriate for identifying early markers of age-associated pathology. For example, in the imaged cohort, thousands of participants are expected to develop Alzheimer’s and Parkinson’s disease by 2030^78^. A notable future QSM resource is the Rhineland Study, which is collecting swMRI phase images in 20,000 individuals at 3T^74^; while QSM has not yet been produced in this study, the complementary age range (≥30 years old) will ultimately enable novel investigations on its own and in combination with UK Biobank data.

Producing accurate and reproduceable χ maps for a dataset at this scale requires a robust, fully automated QSM processing pipeline^16^. We investigated available algorithms for each processing stage to identify an optimal QSM pipeline for the UK Biobank swMRI data (Supplementary Material §1.1). This includes a novel algorithm to automatically detect and exclude voxels with extremely large phase variance, a major source of imaging artefacts (details provided in Supplementary Material §1.1). In our evaluations, we observed considerable variability of χ estimates across different combinations of background field removal and dipole inversion algorithms. Dissemination of our pipeline will thus be crucial for harmonisation of our IDPs with data acquired in novel settings, such as clinical scanners. This will, for example, enable stratification of patients using classifiers^79^ or nomograms^80^ derived from UK Biobank data. To date, two ongoing COVID-19 brain imaging studies have already adopted our QSM pipeline (C-MORE/PHOSP and COVID-CNS)^81^ to process their brain swMRI data.

Using the two-timepoint data from 1,447 subjects (**Supplementary Fig. S10**), we found high cross-scan correlations ( ! > 0.8 ) with χ in four subcortical regions (putamen, caudate, substantia nigra and pallidum). These correlations are higher than the corresponding T2* IDPs (0.6 < ! < 0.8). These four regions show the highest heritability among all QSM and T2* IDPs (**Fig. 5a**), also representing some of the highest heritability values across all brain IDPs in UK Biobank^7^. Notably, these four regions also have the highest χ among all ROIs, and are reported to contain the highest iron concentrations in the brain^82, 83^. χ has demonstrated a strong, positive linear relationship with iron concentration in post-mortem brain tissue^27^. These observations suggest that in structures where iron is the dominant χ source, QSM provides an accurate, reproduceable proxy for tissue iron levels.

### Phenotypic and genetic associations with QSM

The results described in detail above represent approximately 5% of identified phenotypic associations, leaving a rich set of results to be explored further. Some of these phenotypes have previously-established links to χ measures in the brain (e.g., cognitive scores^84, 85^), but many phenotypes lack an obvious interpretation (e.g., χ in caudate and putamen associated with *use of sun/uv protection*). In addition to the extensive literature looking at QSM in neurological conditions, there is a more limited literature in mental health conditions^86, 87^, such as depression and psychosis. Our results supplement this literature, identifying associations between mental health related risk factors including *seen doctor (general practitioner, GP) for nerves, anxiety, tension or depression*; *risk taking;* and *ever taken cannabis* (**Supplementary Fig. S24**).

We described example associations for 19 of the 89 genetic clusters that have a plausible link between gene function and tissue χ. However, we also observed many associations with genetic variants related to biological functions that are not known to be directly related to tissue χ. This includes immune response, regulation of gene expression and cell function (Supplementary §6). This rich set of associations will require additional studies to understand the genetic architecture of magnetic susceptibility in the brain.

In addition to providing insight into “true” associations, voxel-wise maps can help identify spurious associations. For example, several apparent associations at the IDP level seem to be driven by structural atrophy. While all ROI-based IDPs have the potential to be sensitive to atrophy, QSM represents an extreme of image contrast, with opposite sign of χ in grey and white matter. This causes small errors in grey-white boundaries to be amplified into detectable apparent alterations in χ. For example, QSM in substantia nigra has an apparent association with *total BMD (bone mineral density),* but the voxel-wise map exhibits thin, intense associations with a positive-negative pattern at the grey-white boundary that likely reflects atrophy (**Supplementary Fig. S23a,b**). A similar spatial pattern of opposing positive and negative correlations exists for *body mass index (BMI)* in the primary motor and somatosensory areas (**Supplementary Fig. S23c,d**).

### Iron and calcium in quantitative swMRI

Overall, the most consistent and strong pattern of associations identified in our study relate swMRI to iron, both phenotypically (e.g., haemoglobin content) and genetically (e.g., iron homeostasis). Iron is involved in many fundamental biological processes in the brain including neurotransmitter production, myelin synthesis and metabolic processes^88^. Iron homoeostasis is essential to normal brain function, with elevated iron causing oxidative stress and cellular damage^88^. It remains unclear whether abnormal iron accumulation in the brain is a cause or consequence of neurodegeneration in diseases including Alzheimer’s, Parkinson’s and multiple sclerosis^88^.

We also identified multiple strong associations with variants in genes encoding calcium channels. While calcification has been shown to alter tissue χ (χ in calcified lesions have been validated using CT attenuation values^89^), little is known about the impact of other calcium forms on tissue χ. Calcium is essential for many aspects of cell function, including division, differentiation, migration, and death^59^, as well as neurotransmitter and hormone release. Perturbations in calcium homeostasis, including dysregulation of calcium channel activity, have been reported in many neurodegenerative disorders^90, 91^.

These results suggest that the UK Biobank resource can play a key role in the development of swMRI-based biomarkers of iron and calcium. In particular, UK Biobank has unique value for investigating early, asymptomatic disease in individuals who go on to develop neurological conditions. swMRI measures could provide predictive markers for preventative stratification, treatment monitoring and imaging-based screening.

### T2* and QSM in UK Biobank

Imaging at the population scale is a major endeavour that inevitably requires compromises to achieve throughput. While swMRI scans in research settings often use protocols with many echo times lasting 5+ minutes, the UK Biobank swMRI protocol acquires two echoes in 2.5 minutes. QSM involves a single-parameter fit (χ) that can be calculated from one echo, although we explicitly use the two echoes to perform robust coil combination of phase maps. A minimum of two echoes are required to estimate T2* alongside an intercept parameter, making our T2* estimates more sensitive to noise than protocols with more echoes.

Estimation of T2* is also biased by the presence of macroscopic field gradients, induced by air/tissue interfaces or poor shim quality^25^. This bias is exacerbated by the thick slices used in UK Biobank^25^. If not corrected, this can lead to spurious correlations driven by subject-wise variations in field homogeneity rather than tissue χ (**Supplementary Fig. S15**). We introduced deconfounding of macroscopic field gradient for T2* IDPs (see Supplementary §2). As expected, background field gradients do not correlate with our χ estimates (**Supplementary Fig. S16**) and this confound is not needed for QSM.

Despite these potential shortcomings of swMRI in UK Biobank, our results demonstrate that QSM and T2* contribute unique information, as reflected in distinct patterns of associations. Because diamagnetic and paramagnetic constituents manifest differently in QSM and T2* data, a combination of QSM and T2* data may be able to disentangle co-occurring changes of tissue iron and myelin content in neurodegeneration^92^. QSM processing is thus expected to be a valuable addition to the UK Biobank brain imaging resource.

## Methods

### MRI Data acquisition and participant information

We used data from 35,885 participants in the UK Biobank early-2020 release who had susceptibility-weighted MRI (swMRI) data collected. Participants were 53.11% female and aged 45-82yo (64.04±7.5yo) at time of imaging. Of these participants, 1,447 were recruited for a repeat scan approximately 2 years (2.25±0.12y) after the first imaging session. A detailed overview of the neuroimaging acquisition protocols used in UK Biobank brain imaging has been previously described^1^.

Susceptibility-weighted MRI scans were acquired on 3T Siemens Skyra MRI scanners (software platform VD13) with 32-channel head receive coils. swMRI data were acquired using a three-dimensional (3D) dual-echo gradient echo (GRE) sequence with the following parameters: voxel size = 0.8 × 0.8 × 3 mm^3^, matrix size = 256 × 288 × 48 (whole-brain coverage), echo times (TE1/TE2) = 9.4/20 ms, repetition time (TR) = 27 ms and in-plane acceleration = 2, total scan time = 2:34 min. Magnitude and phase data from each receive channel were saved for off-line coil combination, described below.

The UK Biobank brain imaging protocol includes T1- and T2-weighted structural acquisitions that are used in our processing pipeline^15^. Specifically, the T1-weighted structural scan is used to align subjects into a standard-space atlas for the definition of ROIs and other masks, and T2 FLAIR data is used to generate ventricle masks used in QSM referencing. Specifically, brain region masks and image registration references were derived from the T1-weighted structural scans acquired using a 3D MPRAGE protocol (voxel size = 1×1 × 1 mm^3^, matrix size = 208 × 256 × 256, inversion time (TI)/TR = 880/2000 ms, in-plane acceleration = 2, total scan time = 4:54 min)^15^. Ventricle and white matter hyperintensity masks were derived from T2-weighted fluid attenuated inversion recovery (FLAIR) scans (3D SPACE, voxel size = 1.05 × 1 × 1 mm^3^, matrix size = 192 × 256 × 56, TI/TR = 1800/5000 ms, in-plane acceleration = 2, total scan time = 5:52 min)^15^. Both T1 and T2 weighted structural data were processed using the UK Biobank image processing pipeline^15^.

The quality control (QC) pipeline developed for UK Biobank brain imaging^15^ was applied to all subjects included in this study. Several QC measures related to T1-weighted, T2 FLAIR and swMRI data were used here. For example, subjects were excluded if registration from T1 space to standard space failed or had unusable T2 FLAIR data, for example due to excessive head motion, atypical structure and/or anatomical abnormalities. Full details of the quality control pipeline have been previously described^15^. Finally, 35,273 participants were selected in this study whose data were deemed suitable for quantitative susceptibility mapping analysis based on QC measures.

### Quantitative susceptibility mapping processing pipeline

Quantitative susceptibility mapping (QSM) consists of several steps including combination of phase data from individual channels, unwrapping of channel-combined phase data, removal of macroscopic (‘background’) field inhomogeneity, and estimation of voxel-wise χ through dipole inversion (**Fig. 1a**). For each step, many different algorithms have been proposed^16, 93, 94^. To ensure the robustness of our QSM pipeline and select the optimal pipeline for the UK Biobank protocol, we carried out extensive evaluations of established algorithms for each step of the QSM pipeline on a subset of approximately 1,500 subjects^20–24, 95–99^. These evaluations used both quantitative and qualitative metrics, including the presence of streaking artifacts, the observation of large-scale field inhomogeneities on χ maps and cross-subject consistency of both χ maps (in standard space) and χ values in regions of interest (such as subcortical structures). Details of our QSM processing pipeline evaluations are described in Supplementary §1.1.

QSM provides a measure of relative, rather than absolute χ^94^. The presence of an unknown offset in the estimated map for any individual is problematic for comparison of χ values across subjects. To address this, χ values are commonly reported as relative offset with respect to an internal reference region^94, 100^. In this study, we compared the use of three widely-used reference regions: mean χ across (i) the whole brain, (ii) cerebrospinal fluid (CSF) and (iii) a white matter region (forceps minor). Evaluations were performed using subjects who had undergone scanning at two time points (1,447 subjects in total). Specifically, we calculated cross-scan correlations for each of the subcortical QSM IDPs (referenced to these three different regions) as well as subcortical T2* IDPs, with the assumption that negligible changes in χ/T2* occurred during the two timepoints. CSF-referenced QSM IDPs showed the highest consistency (correlation r) across two time points (**Supplementary Fig. S10a**) and CSF in the lateral ventricles was therefore chosen as the reference region. Details of the evaluation of three reference regions are described in the Supplementary §1.2. The final QSM processing pipeline for UK Biobank swMRI data is as follows:

In order to generate a map of the image phase, we first need to combine (average) phase images from individual coil channels. The challenge is that each coil channel has a different unknown phase offset that will lead to phase cancellation (and resulting artefacts) if it is not first removed. Our pipeline combines phase images across channels using the MCPC-3D-S approach^20^. Specifically, a channel-combined phase image was first estimated using the phase difference method (via complex division of two echoes’ phase data, yielding an equivalent TE = TE1 – TE2), masked using the brain mask provided by the UK Biobank image processing pipeline^15^, and then unwrapped using PRELUDE^101^. This unwrapped phase image was scaled to correspond to TE = TE1 and then subtracted from each channel’s first echo phase image respectively to estimate the phase offset for each channel. The estimated phase offsets were subsequently smoothed in the complex domain and subtracted from the phase images (both echoes) of all channels prior to combination. Using this approach, a channel-combined phase image free from any phase cancellation artifacts was generated for each echo.

The brain masks used in this study were based on those used in the standard UK Biobank image processing pipeline^15^. These masks were refined for QSM to detect and remove voxels in the vicinity of sinus cavities. These regions have extremely strong field variations due to the air/tissue interface that induce artifacts in the χ maps. For each echo, we generated a phase reliability map as follows. Channel-combined phase images were converted to complex data (assuming unit magnitude values) and convolved with a 3D spherical kernel (2mm in radius with probabilistic values for voxel locations inside the kernel) for voxels within the brain mask

(only voxels within the intersection of the kernel and brain mask were included for convolution). Our phase reliability map was then derived by calculating the magnitude of the convolved complex data: regions of strong phase variation were indicated by low magnitude values due to phase cancellation within the convolution kernel, while regions with relatively homogeneous phase had magnitude values close to one. Here, the refined mask excluded voxels with phase reliability values lower than 0.6 for the first echo, and 0.5 for the second echo. An additional step was then applied to the refined brain mask to fill any isolated holes (in 3D) in the middle of the bran that were not connected to the sinus cavities using the MATLAB function “imfill”.

The channel-combined phase images were unwrapped using a Laplacian-based algorithm provided by the STI Suite toolbox^21^. The unwrapped phase images were subsequently combined into a single phase image via a weighted sum^22^ over the two echoes to increase signal-to-noise, with weighting factor 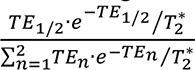 ^22^ (T*_2_ was set as 40 ms for all subjects).

This effectively weights each echo by its predicted signal-to-noise ratio.

The echo-combined phase image was then filtered to remove the background field contributions. Our pipeline uses the variable-kernel sophisticated harmonic artifact reduction for phase data (V-SHARP) algorithm^23^ in the STI Suite toolbox with a maximum kernel size of 12 mm. The phase reliability and hole-filling steps described above were repeated on the brain mask output by V-SHARP, excluding voxels that had phase reliability values lower than 0.7 for the first echo or 0.6 for the second echo (empirically determined). The χ map was generated from the V-SHARP filtered phase data using the refined brain mask. Dipole inversion was performed using the iLSQR algorithm^24^ in the STI Suite toolbox.

The final step is to calculate a reference χ value for each subject to be subtracted from the output χ maps. As described above, we used the χ of CSF in the lateral ventricles as our reference. The χ map was transformed to subject’s native T1 space where the UK Biobank pipeline has previously generated a ventricle mask with BIANCA^15, 102^. Inspection of the χ distribution in the ventricle masks (eroded by a 1mm radius spherical kernel) revealed a bimodal (Gaussian plus inverse gamma) distribution of χ estimates (**Supplementary Fig. S9**). This likely corresponds to both CSF and choroid plexus compartments. To extract the CSF component of this distribution, we used an in-house mixture modelling algorithm^103^. The mean value of the central Gaussian distribution was considered to represent CSF voxels, and was used as the χ reference. The χ map in the original swMRI space was referenced to CSF by subtracting this reference value.

### Image-derived phenotypes generation

In this study, QSM-based image-derived phenotypes (IDPs) were generated relating to 8 subcortical structures (accumbens, amygdala, caudate, hippocampus, pallidum, putamen, substantia nigra and thalamus, left and right) and white matter hyperintensity (WMH) lesions. Masks for each subcortical structure in T1 space (excluding the substantia nigra) have been previously generated using FIRST (FMRIB’s Integrated Registration and Segmentation Tool)^104^ as part of the UK Biobank image processing pipeline. Masks for the substantia nigra used in this study were derived from a substantia nigra atlas^105^ in MNI152 space. Masks for white matter and white matter hyperintensity (WMH) lesions (derived from the BIANCA processing^15, 102^) were provided to us in T1 space from the UK Biobank image processing pipeline, alongside white matter masks generated using FAST (FMRIB’s Automated Segmentation Tool^103^). CSF referenced χ maps were transformed to both subject’s native T1 space and MNI152 space using the provided transformations for each subject from UK Biobank^15^.

FIRST-generated subcortical masks were eroded (2D 3x3 kernel) and used to extract subcortical regions on CSF-referenced χ maps in the T1 space of each subject. The median χ- value (across extracted voxels) was calculated as a separate IDP for each of the left/right subcortical regions. The substantia nigra mask was used to extract the substantia nigra region on CSF referenced χ maps in the MNI152 space of each subject. To account for the morphological variations of substantia nigra across subjects, the substantia nigra mask was refined by excluding voxels that had negative χ values, with median χ values subsequently calculated as a separate IDP for left and right substantia nigra.

For the WMH IDP, the WMH mask was first used to extract the mean χ value in lesions. We then extracted the χ value from normal-appearing white matter, defined as the intersection of FAST- and BIANCA-based white matter mask with the WMH voxels removed. The difference between the estimated χ values of WMH and normal white matter was calculated as an IDP. This approach aims to isolate χ properties that are unique to lesions, as opposed to more global properties of white matter χ.

While most of the QSM IDPs use the same spatial ROIs as existing T2* IDPs, the previous pipelines did not include T2* of substantia nigra or WMH. We thus generated three new T2* IDPs for (substantia nigra left and right, and WMH). This enabled a direct comparison between QSM and T2* for 18 IDPs.

### Outliers and confounds removal

Each IDP’s *N*_*subjects*_ × 1 vector first had outliers removed. Outliers were defined as being greater than 6x the median absolute deviation from the median. The remaining distribution of IDPs was then quantile normalised, resulting in it being Gaussian-distributed with mean zero and standard deviation one.

A recently expanded set of imaging confounds have been proposed in UK Biobank brain imaging^15, 106^ including age, head size, sex, head motion, scanner table position, imaging centre and scan-date-related slow drifts. Unless stated otherwise, the full set of imaging confounds were regressed out from the quantile normalised IDPs before any further analyses. This was crucial to avoid biased or spurious associations between IDPs and other non-imaging measures^1, 106^.

As reported in the literature, T2* estimates are biased by the presence of macroscopic field gradients (for example, induced by air/tissue interface), particularly when imaging voxels are large (in our case, using thick slices)^25^. To reduce such confounding effects on association analyses with UK Biobank T2* IDPs, we first modelled the relationship between macroscopic field gradients and R2* (1/T2*) measurement errors using both simulated and UK Biobank data^25^. This produced a macroscopic field gradient confound variable specific to each subject and brain region, which was generated and regressed out from T2* IDPs. No association was found between the field gradient confounds and QSM IDPs as expected, and thus this particular confound was only applied to T2* data. Details of this additional deconfounding for T2* IDPs are described in the Supplementary §2.

We also observed strong negative correlations between QSM/T2* WMH IDPs and the WMH volume IDP provided by UK Biobank (**Fig. 1d**). This may indicate partial volume effects on the χ and T2* estimates within lesions. QSM and T2* WMH IDPs were thus additionally processed by regressing out the WMH volume.

### Associations between IDPs and non-imaging measures

We investigated 17,485 phenotypes from UK Biobank. Each of the 17,485 phenotypes used here had data from at least 40 subjects. The phenotypes spanned 17 groups of variable types, including early life factors (such as maternal smoking around birth), lifestyle factors (such as diet, alcohol and tobacco consumption), physical body measures (such as BMI, bone density and blood assays), cognitive test scores, health outcomes (such as clinical diagnosis – ICD10 and operative procedures – OPCS4) and mental health variables (such as bipolar and major depression status). These variables were automatically curated using the FUNPACK (the FMRIB UKBiobank Normalisation, Parsing And Cleaning Kit) software to ensure that all phenotypes variables (both quantitative and categorical) were numeric vectors and that resulting correlation coefficients were easy to interpret.

To investigate pairwise associations between IDPs and phenotypes, univariate statistics were carried out using Pearson correlation across all QSM/T2* measures (including 16 subcortical IDPs and 2 WMH IDPs with/without regressing out WMH volume) and 17,485 phenotypes (quantile normalised and fully deconfounded). As UK Biobank phenotypes have varying amounts of missing data, the full set of associations of phenotypes against IDPs had widely-varying degrees-of-freedom. Therefore, it is important to consider *P* values (and not just correlation r); *P* values were calculated and used to identify the strongest associations. Bonferroni multiple comparison correction across the full set of 629,460 (17,485 × 36) correlations was applied, resulting in a -log10*P* threshold of 7.10 (for *Pcorrected* < 0.05). Additionally, a less conservative option for multiple comparison correction is false discovery rate (FDR)^107^, which for a 5% FDR resulted in a -log10*P* threshold of 3.13.

### Associations between IDPs and genetic variants

We carried out genome-wide association studies (GWASs, univariate correlations) for all QSM and T2* IDPs, following a previously-described approach^6, 7^. This was performed using the Spring 2018 UK Biobank release of imputed genetic data. From all subjects with an available χ map, we selected a subset of 29,579 unrelated subjects with recent UK ancestry (to avoid confounding effects from population structure or complex cross-subject covariance). We divided this set into a discovery cohort of 19,720 subjects and a replication cohort of 9,859 subjects. We applied QC filtering to the genetic data including minor allele frequency (MAF) ≥ 0.01, imputation information score ≥ 0.3 and Hardy-Weinberg equilibrium P value ≥ 10^-7^, which resulted in a total of 17,103,079 genetic variants (which are primarily SNPs, single-nucleotide polymorphisms). IDPs had both imaging and genetic confounds regressed out as carried out in Elliott et al.^6^, including the above-described imaging confounds and 40 population genetic principal components (supplied by UK Biobank). IDPs were normalized (resulting in zero mean and unit variance) after the original Gaussianisation and deconfounding. GWAS was performed using the BGENIE software^108^.

Manhattan plots for each of the GWASs were produced, plotting the -log10*P* value for each genetic variant. The standard single-phenotype GWAS threshold (-log10*P* = 7.5)^6, 7^ as well as an additional Bonferroni multiple comparison (accounting for 36 GWASs) corrected threshold

(-log10*P* = 9.06) are shown in the plots. Peak associations with -log10*P* value exceeding 7.5 were extracted and annotated using a method described in Elliott et al.^6^. That is, in a region of high linkage disequilibrium (LD), we only report the genetic variant with the highest association with the IDP because the associations in the local region are most likely all due to a single genetic effect^6, 7, 108^.

After performing the GWAS, we used Peaks software^7^ to automatically generate clusters of peak associations for all IDPs as previously described^6, 7^. A cluster is a set of IDP/variant pairs for which all variants are within a 0.25-cM distance of the IDP/variant pair with highest -log10*P* value in the cluster. Additionally, we used FUMA^43^ to map genetic variants identified in peak associations to related genes and to identify eQTL and chromatin mappings/interactions for these variants.

Following the approach described previously^6^, we examined the heritability (ℎ^(^) of each subcortical IDP using LD score regression (LDSC)^41^ on all available subjects (combining discovery and reproduction cohorts). LD scores were sourced from the European population of the 1000 Genomes Project^109^. Due to limitations in 1000 Genome Project linkage disequilibrium score files, the X chromosome is not considered in this analysis.

### Voxel-wise associations with phenotypes and genetic variants

The MNI152-space versions of CSF referenced χ maps from all subjects were also combined into a 4D MNI152 χ matrix (of size 182 × 218 × 182 × *_*+,-$./*_). Each of the voxel vectors in the 4D MNI152 χ matrix had outliers removed and was quantile normalised and fully deconfounded.

We carried out voxel-wise correlations between the χ maps and non-imaging measures (both phenotypes and genetic variants) that had been identified to have significant associations with QSM-based IDPs. Interrogating χ maps at a voxel-wise level can provide further insight into the spatial localization of associations and can potentially identify additional associated areas which were not captured by the original IDPs, either because a given brain region was explicitly not included or because heterogeneity within a larger brain region diluted an association with a sub-region. Univariate Pearson correlations were performed between fully deconfounded χ data vectors (voxels across the 4D MNI152 χ matrix) and phenotypes or genetic variant data vectors, resulting in 3D correlation maps.

## Acknowledgements

This research was conducted using the UK Biobank Resource under application number 8107. We are grateful to UK Biobank for making the data available and to all UK Biobank study participants, who generously donated their time to make this resource possible. We would like to thank Nicholas Blockley, Stuart Clare, Diego Hernando and Scott Reeder for discussions about deconfounding of macroscopic field gradients; and Saifeng Liu for discussions about phase reliability maps in QSM processing. C.W. is funded, in part, by the China Scholarship Council (CSC). K.L.M., A.B.M.B. and B.C.T. are supported by a Wellcome Trust Senior Research Fellowship 202788/Z/16/Z. S.M.S. and G.D. are supported by a Wellcome Trust Collaborative Award 215573/Z/19/Z. F.A.A. is funded by the UK Medical Research Council and the Wellcome Trust. A.L. is supported by the Horizon 2020 Programme CANDY (Grant No. 847818). The Wellcome Centre for Integrative Neuroimaging is supported by core funding from the Wellcome Trust (203139/Z/16/Z). Computation partially used the Oxford Biomedical Research Computing (BMRC) facility, a joint development between the Wellcome Centre for Human Genetics and the Big Data Institute supported by Health Data Research UK and the NIHR Oxford Biomedical Research Centre.

## Author contributions

C.W ., S.M.S., B.C.T. and K.L.M. designed the research. C.W. developed QSM processing pipeline, created novel IDPs and confounds, and carried out the phenotypic and genetic association analyses. A.B.M.B. interpreted the genetic results. K.L.M., F.A.A. and S.M.S. developed acquisition and core processing pipelines for UK Biobank brain imaging. L.T.E. provided tools for genome-wide associations analysis and carried out heritability analysis. K.L.M., S.M.S., B.C.T, G.D. and J.C.K. evaluated the results. A.L. provided tools for mixture modelling. B.C.T., K.L.M., R.B. and C.F. provided feedback on the QSM processing pipeline. C.W., A.B.M.B., B.C.T. and K.L.M. wrote the manuscript, which was edited by all authors.

## Data availability

All source data (including raw and processed brain imaging data and genetics data) are (or will be) available from UK Biobank via their standard data access procedure (see http://www.ukbiobank.ac.uk/register-apply).

## Code availability

The image processing pipelines of the MRI data in the UK Biobank project can be found at http://www.fmrib.ox.ac.uk/ukbiobank or https://git.fmrib.ox.ac.uk/falmagro/UK_biobank_pipeline_v_1. Custom-written MATLAB code for QSM processing is freely available at https://git.fmrib.ox.ac.uk/cwang/uk_biobank_qsm_pipeline and will be added to the core UK Biobank brain imaging processing pipeline.

The following software packages were used in this work:

- FMRIB Software Library (FSL) v6.0, https://fsl.fmrib.ox.ac.uk/fsl/fslwiki
- STI Suite, https://people.eecs.berkeley.edu/~chunlei.liu/software.html
- MEDI toolbox, http://pre.weill.cornell.edu/mri/pages/qsm.html
- Mixture modelling, https://github.com/allera/One_Dim_Mixture_Models
- Peaks v1.0, novel software for extracting clusters from multi-phenotype GWASs: https://github.com/wnfldchen/peaks
- bgenie v1.3, software for efficient GWASs on high-dimensional phenotype data: https://jmarchini.org/bgenie/
- LDSC v1.0.1, software for heritability analysis from summary statistics (linkage score regression): https://github.com/bulik/ldsc/

