## Supplementary Material for "Phenotypic and genetic associations of quantitative magnetic susceptibility in UK Biobank brain imaging"

### Section 1: QSM pipeline evaluations

#### 1.1 Evaluation of processing algorithms

Quantitative susceptibility mapping (QSM) consists of several steps, including combination of phase data from individual channels, unwrapping of channel-combined phase data, removal of macroscopic ('background') field inhomogeneities, and estimation of voxel-wise magnetic susceptibility ($\chi$) through dipole inversion. For each step, many different algorithms have been proposed. To ensure the robustness of our QSM pipeline and to select the optimal pipeline for the UK Biobank protocol, we carried out extensive evaluations of established algorithms (from widely-used and publicly-available toolboxes) for each step.

*Combination of multi-channel phase data*

Robust combination of multi-channel phase data is crucial for performing QSM. Different channels have different phase offsets ($\varphi_{0,j}$) associated with the respective coil-sensitivity field. If left unaccounted for, these offsets will lead to signal cancelation and open-ended fringe lines in the resulting coil-combined phase maps. As UK Biobank swMRI protocol collects two echoes (TEs = 9.4 and 20 ms) without performing reference scans (for coil sensitivity estimation), only a handful of algorithms can be used for coil combination. In this work, we compared two different coil combination methods: (1) phase difference and (2) MCPC-3D-S[1].

Two-echo phase from the $j^{th}$ coil can be written as:

$$\varphi_{1,j} = \gamma \Delta B \cdot TE_1 + \varphi_{0,j} \quad [1]$$

$$\varphi_{2,j} = \gamma \Delta B \cdot TE_2 + \varphi_{0,j} \quad [2]$$

The phase difference method uses complex division of the two-echo phase to eliminate the time-independent phase offset term, $\varphi_{0,j}$. This enables direct combination with an equivalent TE = $TE_2$-$TE_1$, where:

$$\varphi_{diff,comb} = \sum_j \varphi_{diff,j} = \sum_j \gamma \Delta B \cdot (TE_2 - TE_1) \quad [3]$$

and $\varphi_{diff,comb}$ is the coil-combined phase difference map. This method is straightforward, but reduces the signal-to-noise ratio (SNR) due to the subtraction of two echoes.

MCPC-3D-S is a recently proposed algorithm that uses a scaled coil-combined phase difference map to estimate the phase offset in each coil, defining:

$$\varphi_{0,j,est} = \varphi_{1,j} - unwrap(\varphi_{diff,comb}) \cdot \frac{TE_1}{TE_2 - TE_1} \quad [4]$$

where $unwrap(\varphi_{diff,comb})$ corresponds to an unwrapped phase difference map (required when $TE_2 \neq 2 \cdot TE_1$), and $\varphi_{0,j,est}$ is the estimated phase offset for coil j. $\varphi_{0,j,est}$ estimates are spatially-smoothed and subtracted from $\varphi_{1,j}$ and $\varphi_{2,j}$ to generate individual channel phase maps without offsets. These phase maps can be subsequently coil-combined, resulting in a single phase map for each echo.

Although MCPC-3D-S is slower than the phase difference method (due to multiple additional steps including phase unwrapping, here performed using PRELUDE[2]), it generates coil-combined phase data with higher SNR verses the phase difference method, as shown in **Fig. S1**.

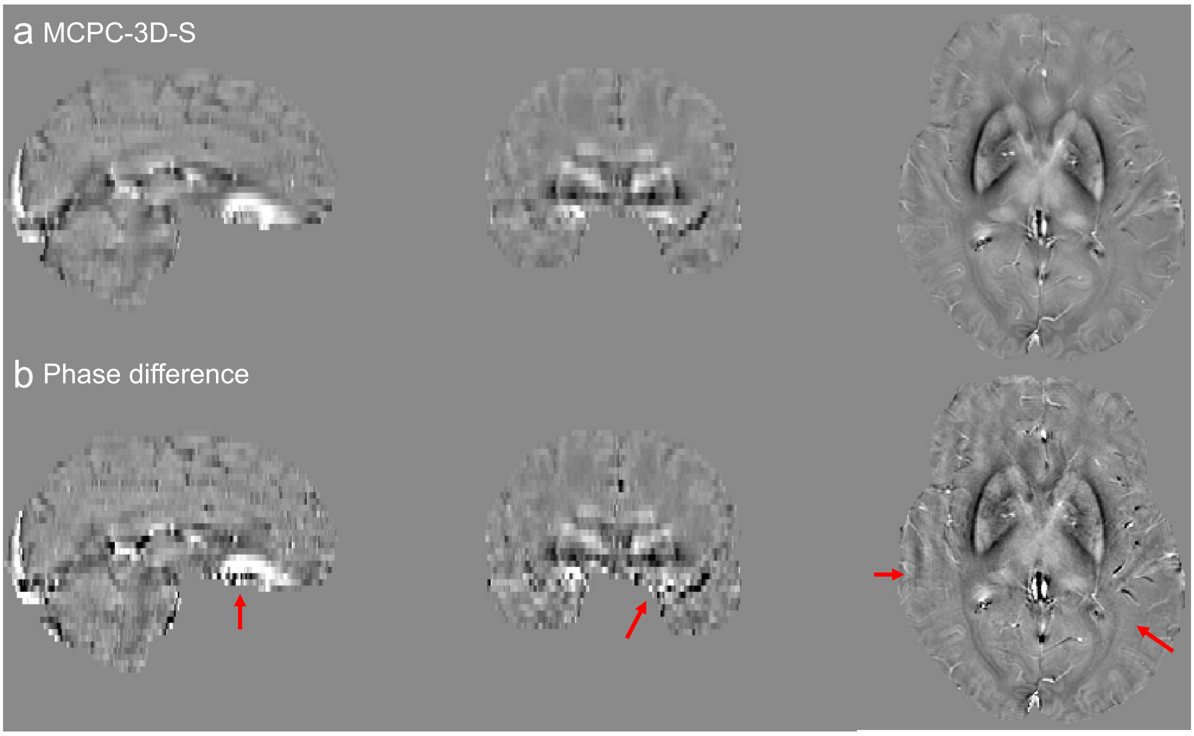


**Figure S1** Example filtered phase data from a single UK Biobank subject using MCPC-3D-S **(a)** and phase difference **(b)** channel combination. Overall, MCPC-3D-S (a) generated images with higher SNR and fewer imaging artefacts (red arrows) than phase difference (b).

*Unwrapping of channel-combined phase*

There are a number of algorithms which have been developed for phase unwrapping. In this study, we compared a path-based (FSL's PRELUDE [2]) and a Laplacian-based[3] (from the STI suite toolbox https://people.eecs.berkeley.edu/~chunlei.liu/software.html) algorithm. Overall, the Laplacian-based algorithm demonstrated rapid phase unwrapping with smoother phase variance in (noisier) regions with large phase jumps verses PRELUDE, as shown in **Fig. S2**. Laplacian-based algorithms perform a degree of background field removal when unwrapping phase maps, which restricts their use when quantitative phase values are required (e.g. the coil-combination step). However, as unwrapped phase data are subsequently filtered in QSM (to remove background field contributions), this does not restrict their use for generating $\chi$ maps.

The unwrapped phase images were subsequently combined into a single phase image via a weighted sum[4] over the two echoes to increase signal-to-noise, with weighting factor $\frac{TE_{1/2} \cdot e^{-TE_{1/2}/T_2^*}}{\sum_{n=1}^{2} TE_n \cdot e^{-TE_n/T_2^*}}$ [4] ($T_2^*$ was set as 40 ms for all subjects). This effectively weights each echo by its predicted signal-to-noise ratio.

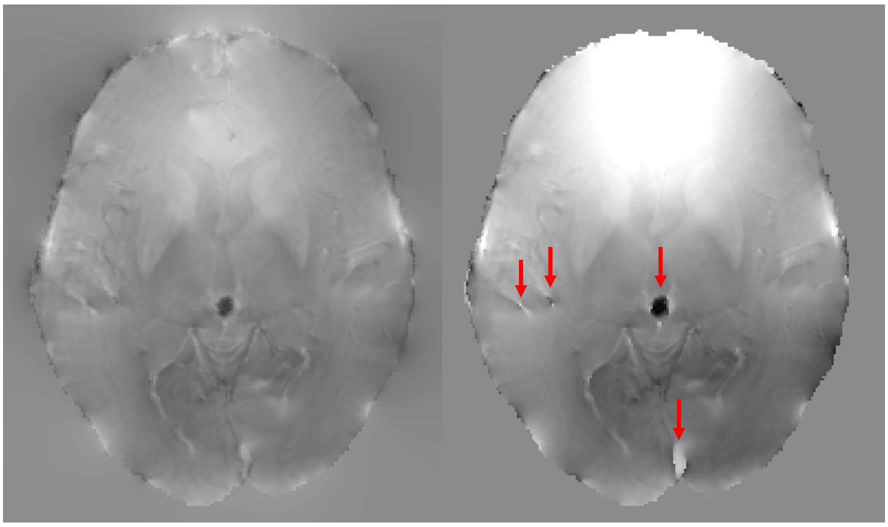


**Figure S2** Unwrapped phase image from an example UK Biobank subject. Red arrows indicate boundaries with large phase jumps in the unwrapped phase map using PRELUDE, which are not present using the Laplacian-based algorithm.

*Phase reliability maps*

Unreliable phase estimates can lead to artefacts and large-scale image inhomogeneities on resulting $\chi$ maps. To address this, a phase reliability map was estimated per subject to identify and remove these voxels. Here, channel-combined phase images were first converted to complex data (assuming unit magnitude values) and convolved with a 3D spherical kernel (2mm radius). In this procedure, the spherical kernel was normalised to account for the fraction of each voxel contained in the kernel, and a further normalisation correction was performed for voxels in close/immediate vicinity to the brain boundary (where only a fraction of the convolution kernel included brain tissue). The phase reliability map was subsequently derived by calculating the magnitude of the convolved complex data: regions of strong phase variation were indicated by low magnitude values due to phase cancellation from the convolution step, while regions with relatively homogeneous phase had magnitude values close to one, as shown in **Fig. S3**. The threshold of the phase reliability map for each echo was (empirically) determined after visual inspection of ~100 subjects (0.6 for first echo and 0.5 for second echo before background field removal; 0.7 for first echo and 0.6 for second echo before dipole inversion), to exclude unreliable voxels (predominantly in the vicinity of sinus cavities). An additional step was then applied to the refined brain mask to fill any isolated holes (in 3D) in the middle of the bran that were not connected to the sinus cavities using the MATLAB function “imfill”.

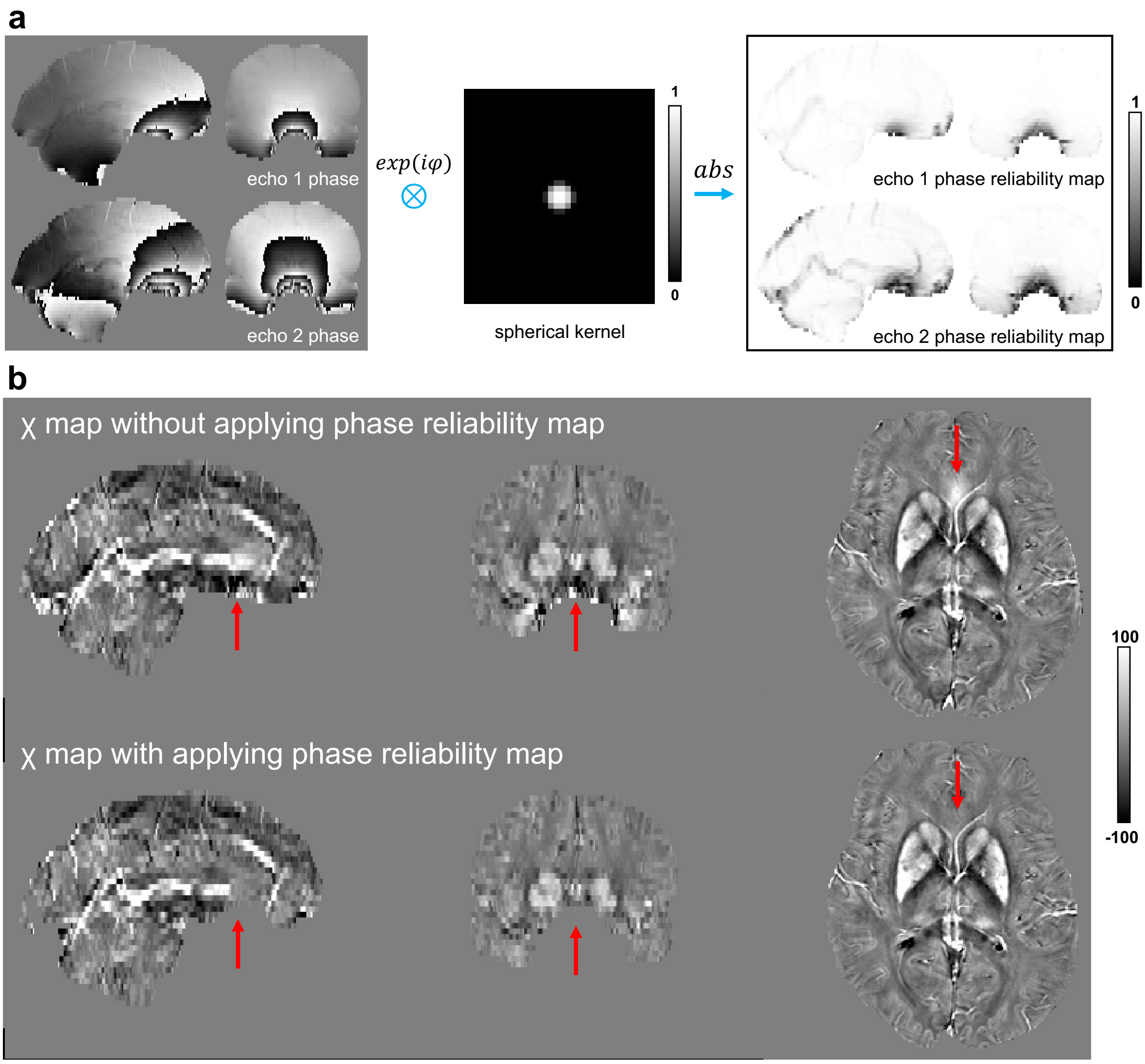


**Figure S3 (a)** Phase reliability map pipeline for each echo in a single UK Biobank subject. ⊗ represents convolution in complex space, where phase data ($\varphi$) were converted to exp $(i \cdot \varphi)$. Phase reliability map was calculated as the magnitude (absolute value) of the convolved complex data **(b)** Example χ map generated without (top) and with (bottom) the phase reliability map correction. Red arrows indicate regions with strong phase variations that induced large field inhomogeneities. The phase reliability correction removes voxels with large phase variations at the brain edge, resulting in χ maps with fewer artefacts.

*Background field removal and dipole inversion*

Removal of background field contributions and subsequent dipole inversion are the final two steps required to generate χ maps, with a number of different algorithms proposed for each step. Here, we evaluated combinations of these algorithms based on the final χ map. These evaluations used both quantitative and qualitative metrics, including the observation of large-scale inhomogeneities and streaking artefacts on χ maps, and cross-subject consistency of both χ maps (in standard space) and χ values in regions of interest (subcortical structures).

We compared 5 different background field removal algorithms (V-SHARP[5], PDF[6], iHARPERELLA[7], iRSHARP[8] and LBV[9]), 3 different dipole inversion algorithms (iLSQR[10], STAR-QSM[11] and MEDI[12]) and 1 single-step background field removal and dipole inversion algorithm (fast TFI[13]) resulting in the evaluation of 16 $(3 \times 5 + 1)$ different combinations.

Background field removal:

Here we present evaluations using group averaged $\chi$ maps over the 50 subjects for the different background field removal algorithms. Although evaluations were performed using all of the proposed dipole inversion algorithms, for conciseness here we display results using iLSQR only. Details of our dipole inversion algorithm evaluations are provided in the following section.

Figure S4 compares V-SHARP with PDF, iHARPERELLA, iRSHARP, and LBV. Overall, we found that V-SHARP provided the best performance, generating $\chi$ maps without observable large-scale inhomogeneities and a low cross-subject standard deviation on resulting $\chi$ maps.

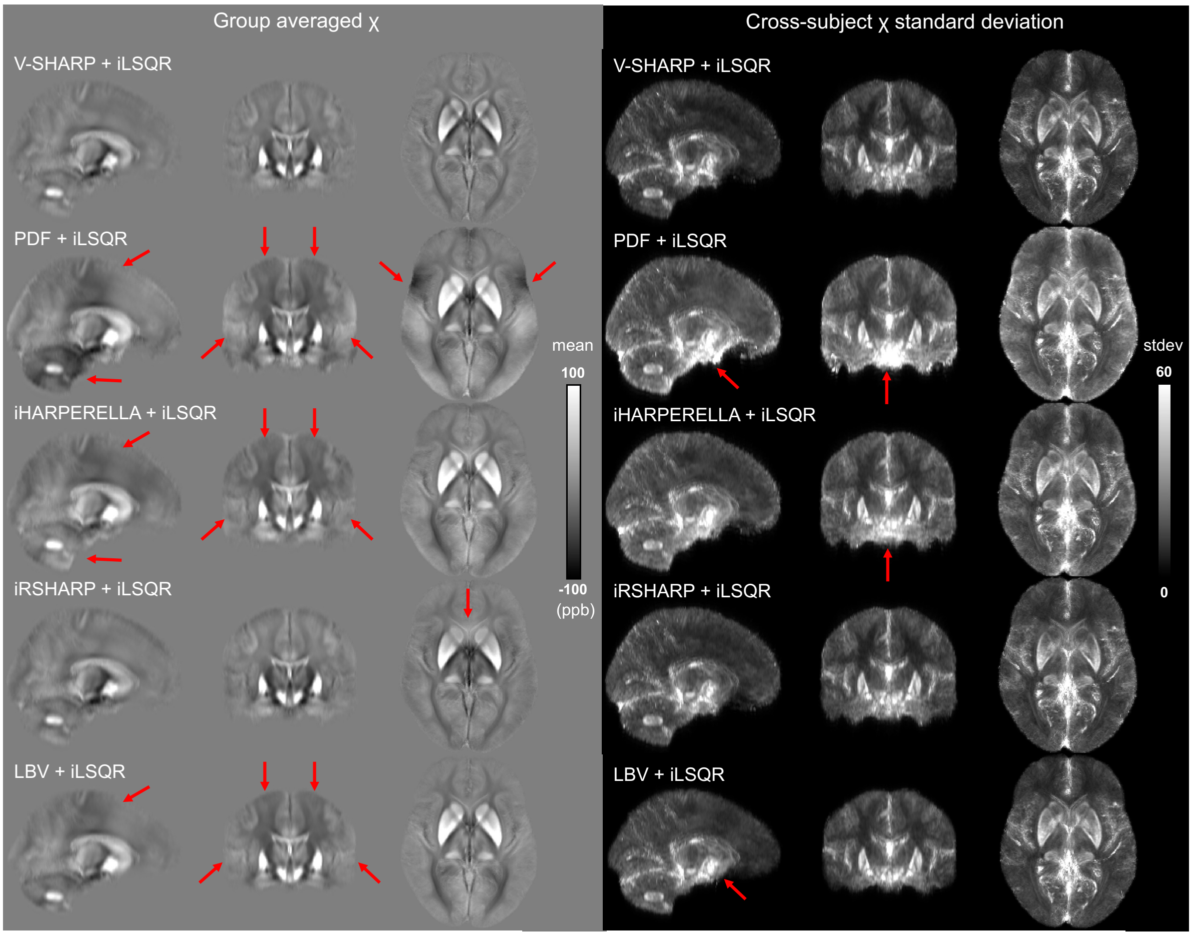


**Figure S4** Group averaged $\chi$ maps (left) and standard deviation (stdev) maps (right) from 50 subjects. $\chi$ maps were produced using V-SHARP/PDF/iHARPERELLA/iRSHARP/LBV and iLSQR. Red arrows highlight regions of large-scale inhomogeneity (consistent across subjects) and cross-subject variation on resulting $\chi$ maps when compared to V-SHARP and iLSQR. Although iRSHARP and V-SHARP are derived from the same technique (SHARP), for UK Biobank data the V-SHARP implementation was found to remove more field inhomogeneities in brain regions in close vicinity to our regions of interest, including the caudate, putamen and pallidum.

Dipole inversion algorithms:

Here we present evaluations using group averaged $\chi$ maps over the 50 subjects for the different dipole inversion algorithms. Although evaluations were performed using all of the proposed background field removal algorithms, again for conciseness here we display results using V-SHARP only, which was the method adopted for background field removal (**Fig. S4**). Comparisons between V-SHARP + iLSQR, STAR-QSM and MEDI, in addition to the single-step fast TFI algorithm are shown in **Fig. S5**.

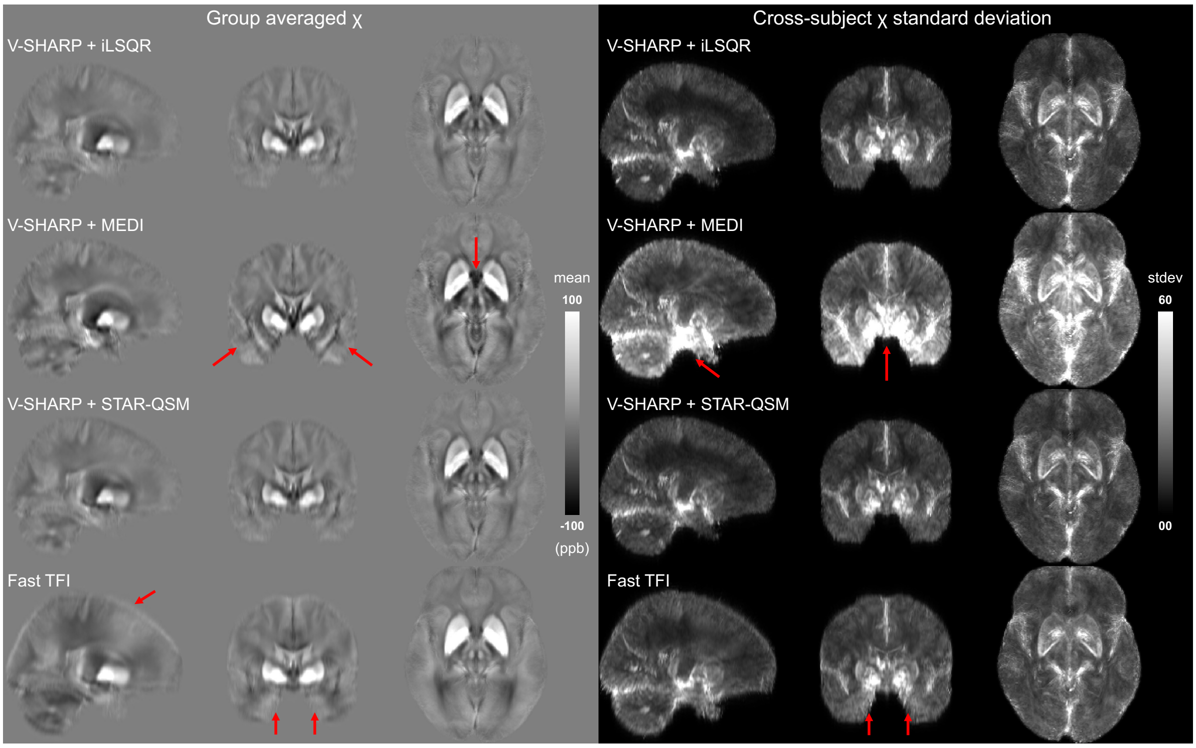


**Figure S5** Group averaged χ maps (left) and standard deviation (stdev) maps (right) from 50 subjects. χ maps were produced using V-SHARP and iLSQR/STAR-QSM/MEDI, in addition to the single-step fast TFI algorithm. Red arrows highlight regions of large-scale inhomogeneity (consistent across subjects) and cross-subject variation on resulting χ when compared to V-SHARP and iLSQR. The group averaged χ maps using iLSQR and STAR-QSM visually appeared of very similar quality.

STAR-QSM and iLSQR produced similar appearing χ maps. We therefore carried out an expanded comparison using phase data without the phase reliability map correction, to simulate the circumstances where voxels with large field variations remained in the dataset. **Fig. S6** demonstrates that iLSQR outperformed STAR-QSM in such a situation.

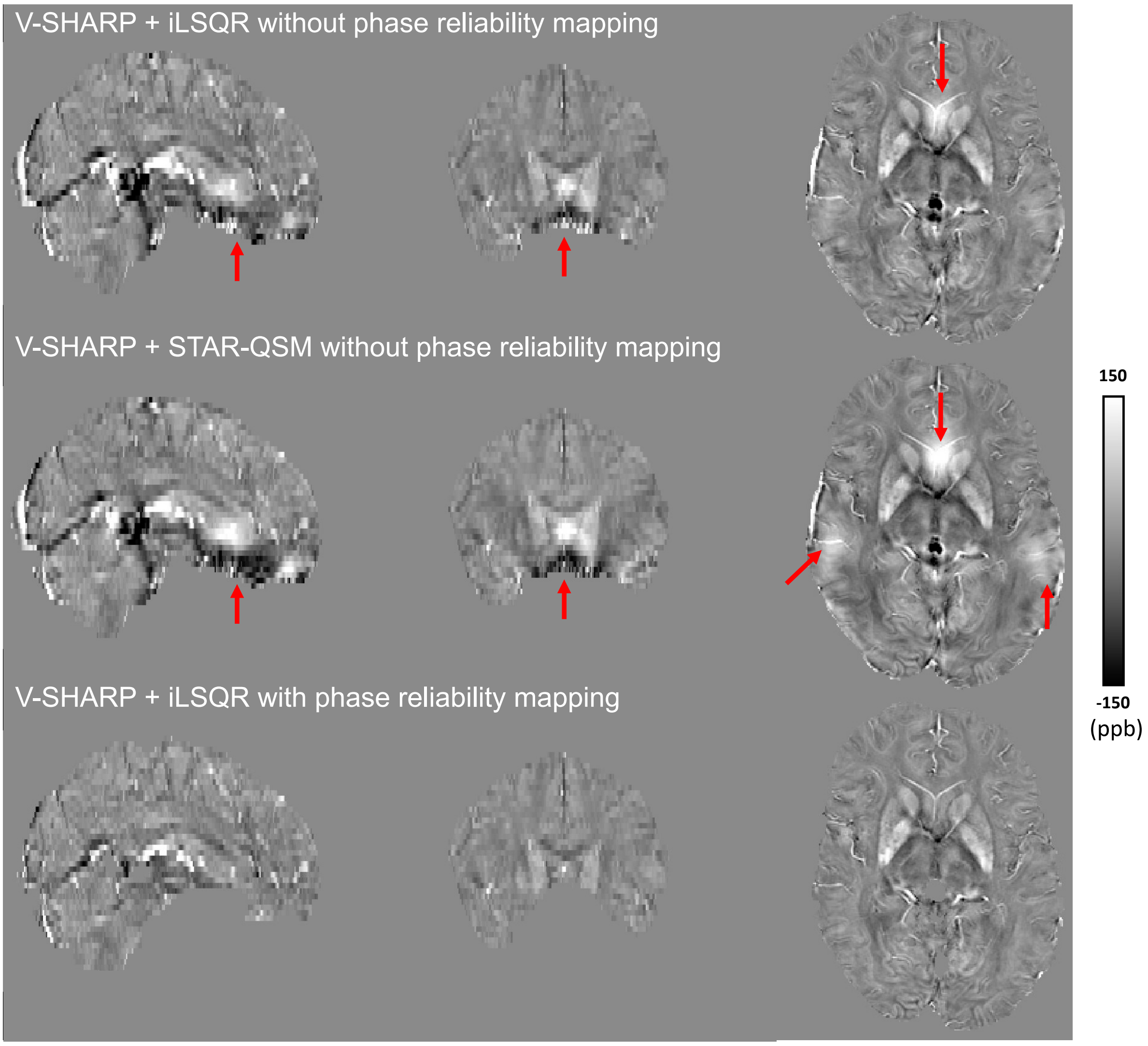


**Figure S6** Example χ map from a single UK Biobank subject using V-SHARP and iLSQR/STAR-QSM without the phase reliability map correction (**Fig S3**). Overall, iLSQR demonstrated improved suppression of streaking artefacts and image inhomogeneities on resulting χ maps (top) verses STAR-QSM (middle), important if the phase reliability correction failed to remove all voxels with unreliable phase information. Remaining V-SHARP and iLSQR image inhomogeneities are subsequently eliminated with the phase reliability correction (bottom).

Finally, we evaluated the different background field and dipole inversion algorithms by calculating the median χ in 14 subcortical ROIs (thalamus, caudate, putamen, pallidum, hippocampus, amygdala, accumbens, left and right separated) (**Fig. S7**). Although different combinations of algorithms showed varying degrees of artefacts on χ maps (**Figs. S4 and S5**), median χ measures in the subcortical ROIs are broadly in similar ranges. The combination of V-SHARP and iLSQR generally showed the smallest within-ROI variance.

Our comparisons determined that the combination of MCPC-3D-S, Laplacian-based phase unwrapping, masking with the phase reliability correction, V-SHARP and iLSQR was the optimal pipeline for UK Biobank swMRI data.

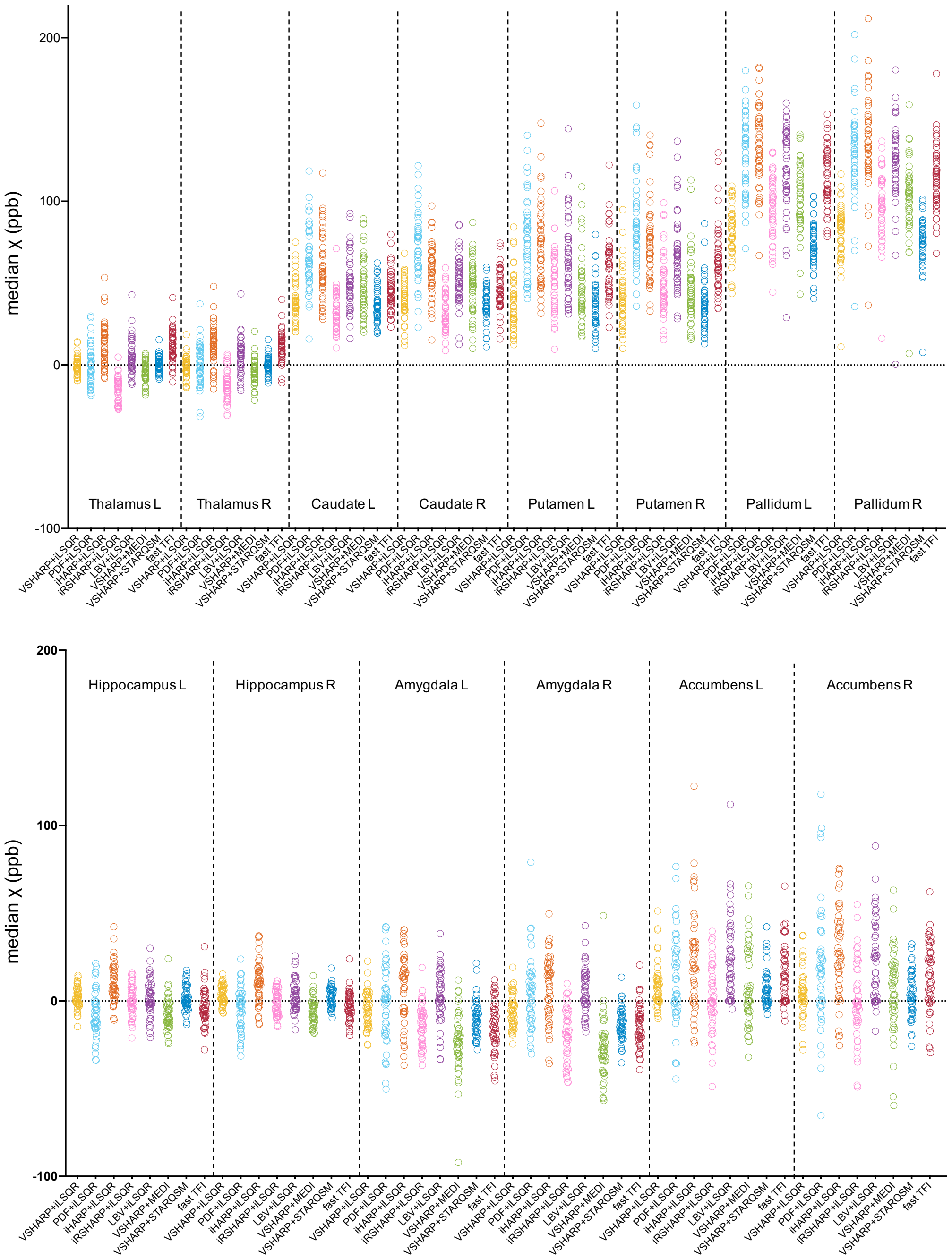


**Figure S7** Comparison of median χ values derived from χ maps for 8 combinations of algorithms. Here, each point represents a single χ estimate from each of the 50 subjects. For each ROI, different combinations generated median χ values covering a broadly similar range. The combination of V-SHARP and iLSQR generally showed the smallest within-ROI variance.

### 1.2 evaluations of reference region for χ maps

QSM provides a measure of relative, rather than absolute χ[14]. The presence of an unknown offset in the estimated map for any individual is problematic for comparison of χ values across subjects. To address this, χ values are commonly reported as offset with respect to an internal reference region. In this study, we compared the use of three widely-used reference regions (**Fig. S8**): mean χ across (i) the whole brain, (ii) cerebrospinal fluid (CSF) and (iii) a white matter region (forceps minor).

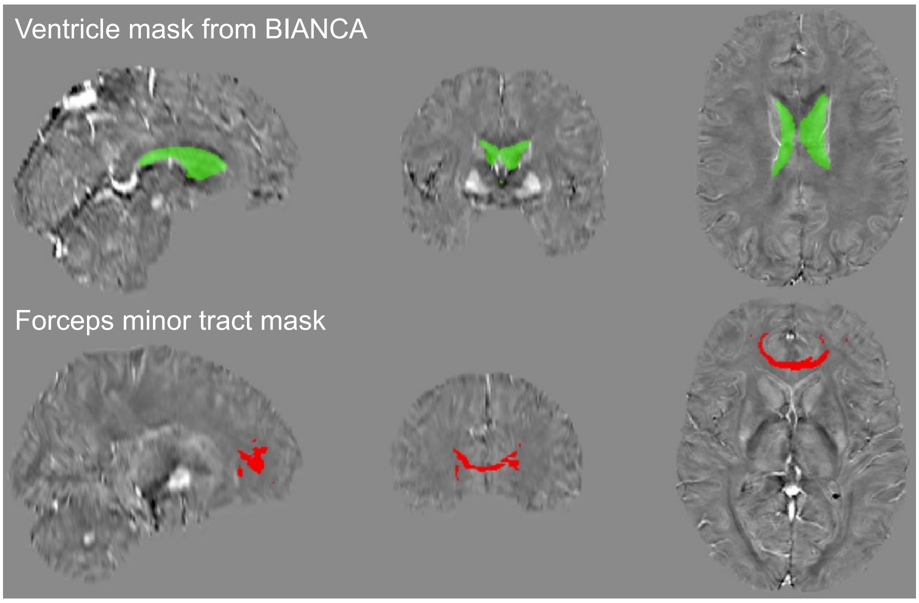


**Figure S8** Ventricle and forceps minor tract masks used for the CSF and white matter reference, overlayed on an example χ map (single subject). Ventricle masks for each subject were previously generated as part of UK Biobank, extracted from the T2 FLAIR data using BIANCA[15]. The forceps minor tract mask was derived from the diffusion MRI data using AutoPtx[16].

The χ distribution in the ventricle mask used for CSF referencing (**Fig. S9**) contained a bimodal (Gaussian and inverse gamma) distribution, which likely corresponds to CSF and the choroid plexus. To extract the CSF component of this distribution, we used an in-house mixture modelling algorithm[17]. The mean of the central Gaussian distribution was considered to represent CSF voxels, and was used as the CSF χ reference.

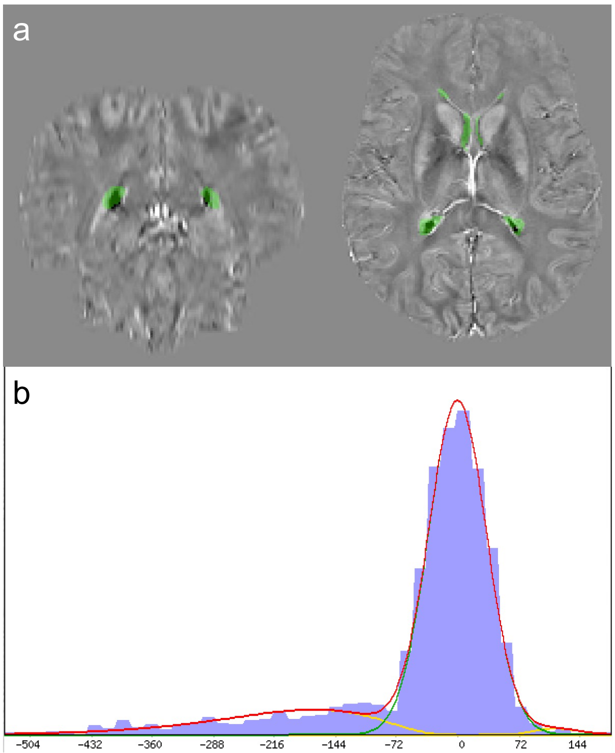


**Figure S9 (a)** Example ventricle mask overlaid on top of a χ map for a single subject. **(b)** distribution of χ in the ventricle mask. The negative voxels likely correspond to the choroid plexus, resulting in the left "tail" (negative inverse gamma) of the χ distribution. The CSF reference χ estimate was measured as the mean of the main Gaussian distribution.

Evaluation of the three reference regions were performed using UK Biobank subjects who had undergone scanning at two time points (1,447 subjects in total) with second imaging session performed approximately 2 years (2.25±0.12y) after the first imaging session. Specifically, we compared cross-scan consistency of $\chi$ maps referenced to these three different regions (**Fig. S10**) with the assumption that negligible changes in $\chi$ occurred during the two timepoints.

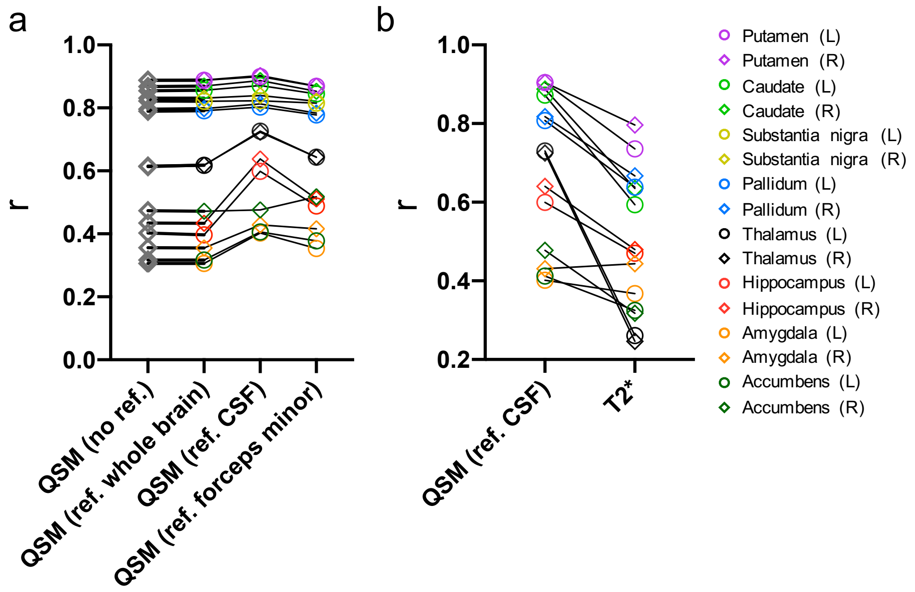


**Figure S10** Correlation between QSM IDPs obtained at the first and second imaging visit. (a) Overall, the CSF referenced IDPs yielded the highest r values (most evident in the thalamus and hippocampus), indicating that CSF is the most robust reference among the three regions. (b) QSM IDPs typically display higher reproducibility verses the T2* IDPs.

### Section 2: Additional deconfounding for T2* IDPs

Estimation of a biologically-meaningful T2* reflecting cellular compartments is confounded by the presence of macroscopic field gradients induced by air/tissue interfaces or poor magnetic field shim quality[18]. In large-cohort epidemiological studies, this could lead to spurious correlations driven by subject-wise variations in field homogeneity (for example, geometry of the sinuses, or other anatomical structures outside the brain), rather than cellular phenomena. In the presence of macroscopic field gradients, the gradient-echo signal can be modelled as[18]:

$$\mathrm{S}(\mathrm{TE}, r_0) = \int_{R^3} S_0\, e^{-R2^*(r)TE} e^{j2\pi\gamma\Delta B_0(r)TE} \cdot SRF(r - r_0)dr \qquad [5]$$

where $\mathrm{S}(\mathrm{TE}, r_0)$ is the measured signal at echo time TE and location $r_0$, $R2^*(r)$ is the spatial distribution of $R2^*$ ($= 1/T2^*$), $\gamma\Delta B_0(r)$ is the magnetic field offset (in Hz), and $SRF(r - r_0)$ is the spatial response function (SRF) of a voxel centered at $r_0$. As spatial resolution along the slice direction (z-dimension) is typically lower than the in-plane dimensions, Eq. [5] can be simplified as[18]:

$$\mathrm{S}(\mathrm{TE}, z) = \int_{R} S_0\, e^{-R2^*(z)TE} e^{j2\pi\gamma\Delta B_0(z)TE} \cdot SRF(z - z_0)dz \qquad [6]$$

Here, we describe our approach to estimate and deconfound for the impact of macroscopic field gradients on the UK Biobank T2* analysis.

Simulated impact of macroscopic field gradients – 2D and 3D acquisitions:

Simulations are based on the procedure by Hernando et al.[17]. Assuming a linear through-slice field variation ($\Delta B_0(z) = \Delta B_0(z_0) + G[z - z_0]$), for a 2D acquisition the SRF is a boxcar function (**Fig. S11a**, first row – left). This leads to:

$$\mathrm{S}(\mathrm{TE}, z) = S_0 e^{-R2^{*}TE} e^{j2\pi\gamma\Delta B_0(z_0)TE} \cdot \Delta z \cdot sinc(\gamma G_z \Delta z TE) \quad [7]$$

where $G_z$ is the macroscopic field gradient along the slice direction and $\Delta z$ is the slice thickness. For 2D acquisitions with an ideal slice profile, the measured signal is thus modulated by $sinc(\gamma G\Delta zTE)$ (**Fig. S11a**, first row - right), with larger field gradients (G) or thicker slices ($\Delta z$) leading to faster signal decay. For 3D acquisitions, the SRF is a sinc-like function (**Fig. S11a**, second row - left), leading to a measured signal modulated by a boxcar-like function (**Fig. S11a**, second row - right).

Simulated impact of macroscopic field gradients – UK Biobank swMRI protocol:

The UK Biobank swMRI protocol is a 3D sequence with 3 mm slice thickness, incorporating k-space filtering (windowing) to remove Gibbs ringing and improve SNR. This leads to an SRF consisting of a rapidly-decaying sinc-like function (**Fig. S11a**, third row - left), with the measured signal modulated by a smoothed boxcar-like function (**Fig. S11a**, third row - right).

To estimate the bias on T2* estimates on UK Biobank swMRI data, we first simulated the impact of macroscopic field gradients using the UK Biobank swMRI protocol (**Fig. S11a**, third row). We subsequently used these simulations to model the relationship between estimated R2* ($s^{-1}$) (as demonstrated by Eq. [5-7], calculation in R2* is more straightforward than in T2*) and Gz (Hz/mm) (**Fig. S11b**), using this relationship to estimate parameters to inform the UK Biobank T2* deconfounding (**Fig. S11c**).

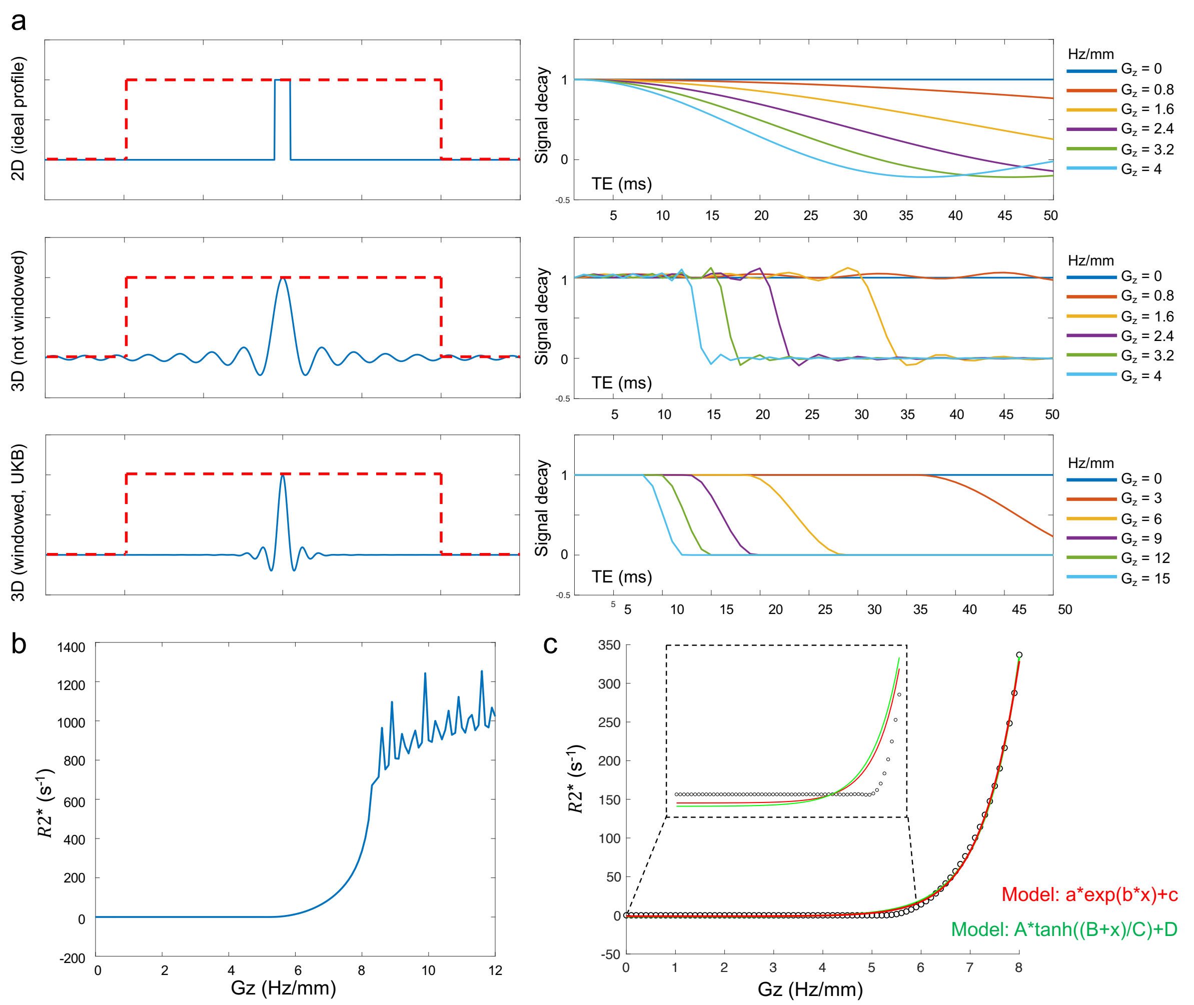


**Figure S11 (a)** Simulated signal modulation in the presence of a linearly varying macroscopic field gradient (Gz), based on the procedure proposed by Hernando et al.[18]. On the left, the signal amplitude is indicated as a boxcar function (red line, representing a large homogeneous object), and the SRF (blue line) is displayed for 3 different imaging scenarios: 2D (10 mm slice – first row), 3D (10 mm slice – second row), and the UK Biobank swMRI protocol (3D, 3mm slice with k-space windowing – third row). One the right, the corresponding signal modulation is displayed as a function of TE and Gz. The presence of macroscopic field gradients (Gz > 0) leads to faster signal decay. **(b)** Simulated R2* estimates based on the UK Biobank protocol, as a function of Gz. The R2* estimate is dominated by noise when Gz > 8 Hz/mm, arising due to the negligible signal available for the second UK Biobank swMRI echo (20ms). **(c)** Two models were used to fit to the R2*-Gz curve over the range [0 8] Hz/m: *a*exp(b*x)+c* and *A*tanh((B+x)/C)+D*. Both models yielded similar fitted curves with $R^2 > 0.99$. Here, a = 0.0052, b = 1.384, c = -1.6924; A= 922.45, B = -9.0029, C = 1.3161, D = 921.23.

Experimental macroscopic field gradient deconfounding:

To reduce the confounding effect of field gradients on the association analyses with T2* IDPs, we generated macroscopic field gradient maps along the slice-direction (**Fig. S12**) for all UK Biobank subjects. Median gradient magnitude values were subsequently calculated in each subcortical ROI as a summary measure of the background field gradient, used to develop an additional set of confounds when performing our association analysis. The distribution of median gradient magnitude (Gmag) for subcortical ROIs is shown in **Fig. S13.**

To generate the macroscopic field gradient maps, the two-echo coil-combined phase data from each subject was unwrapped using PRELUDE and averaged to generate $\gamma\Delta B_{averaged}$. PRELUDE was chosen as it does not remove any background field components (which could bias field gradient estimates). $\gamma\Delta B_{averaged}$ was subsequently filtered using V-SHARP, $\gamma\Delta B_{V\_SHARP}$, with the macroscopic field of each subject estimated as:

$$\gamma\Delta B_{background} = \gamma\Delta B_{averaged} - \gamma\Delta B_{V\_SHARP} \quad [8]$$

Gmag was subsequently generated by taking the gradient magnitude (along the z-direction) of $\gamma\Delta B_{background}$ (**Fig. S12**).

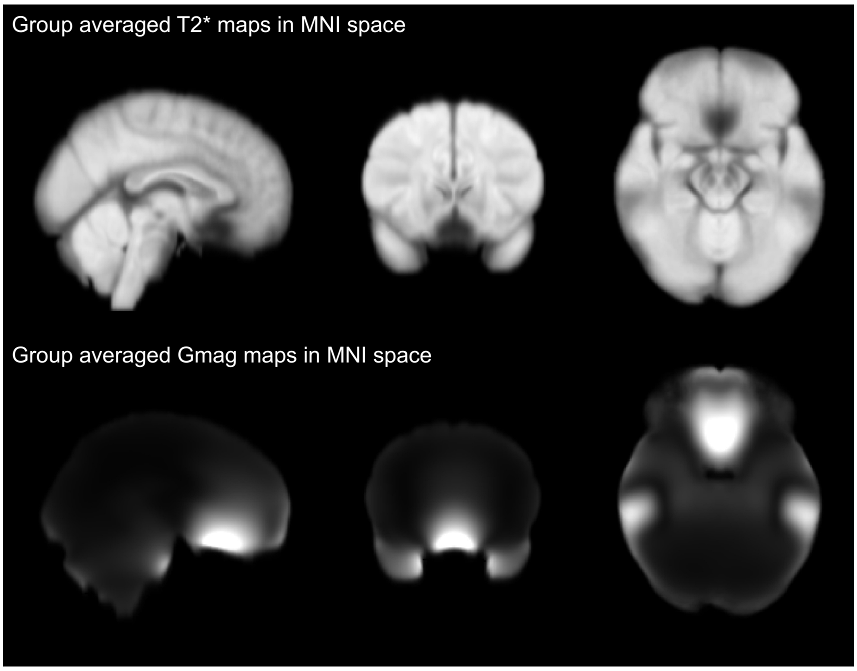


**Figure S12** Group average T2* map (top) and Gmag (bottom) over 200 subjects. The sinus cavity and ear canals are the major sources of variations in background fields due to air/tissue interfaces.

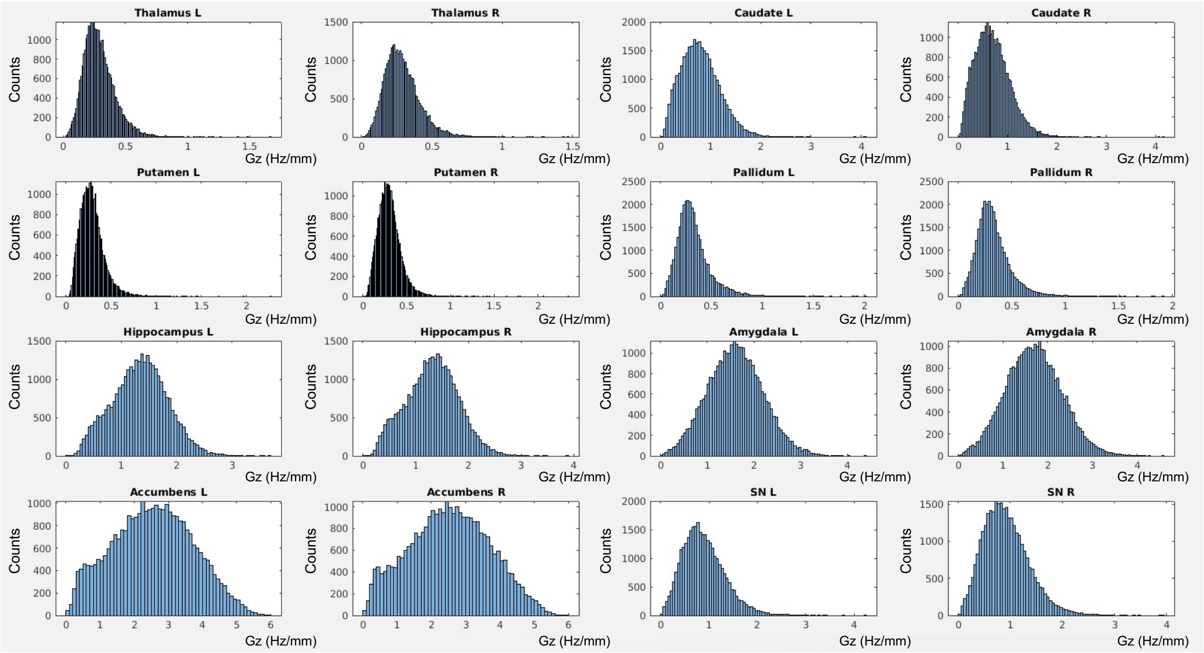

**Figure S13** Histograms of median Gmag in 16 ROIs across all UK Biobank subjects. Only the accumbens, amygdala and hippocampus regions contain a wide Gmag distribution exceeding 2 Hz/mm.

We subsequently modelled the voxelwise relationship between macroscopic field gradients (**Fig. S12**) and R2* (1/T2*) (**Fig. S14**), to produce a set of confound to account for the background field gradient. To achieve this, we evaluated a series of different models (based on the simulations in **Fig. S11c**) to identify the relationship between R2* and Gmag, as shown in **Fig. S14**. The tanh model (*A*tanh((B+x)/C)+D*) model showed the best performance, defining *B*=-6.06 and *C*=1.87, similar to the fitting parameters estimated using the simulated data (**Fig. S11c**, *B* = -9.00, *C* = 1.32). We used these fitting parameters establish a linear relationship between the R2* and median Gmag measures, setting $Gmag_{deconf}$ = *tanh((-6.11+Gmag)/1.87)*.

Changes in the voxelwise R2* estimates manifest in regions with large macroscopic field gradients (**Fig. S14**). From the median subcortical ROI analysis (**Fig. S13**), only the accumbens, amygdala and hippocampus are confounded by large macroscopic field gradients (Gmag > 2 Hz/mm). Therefore, we only performed deconfounding (linear regression) for T2* IDPs in the accumbens, amygdala and hippocampus.

As shown in **Fig. S15**, spurious associations between T2* accumbens IDPs and sinus/nasal-related phenotypes (including ICD10 codes related to *Nasal polyp* and *Chronic sinusitis*) are dramatically reduced after deconfounding for macroscopic field gradients.

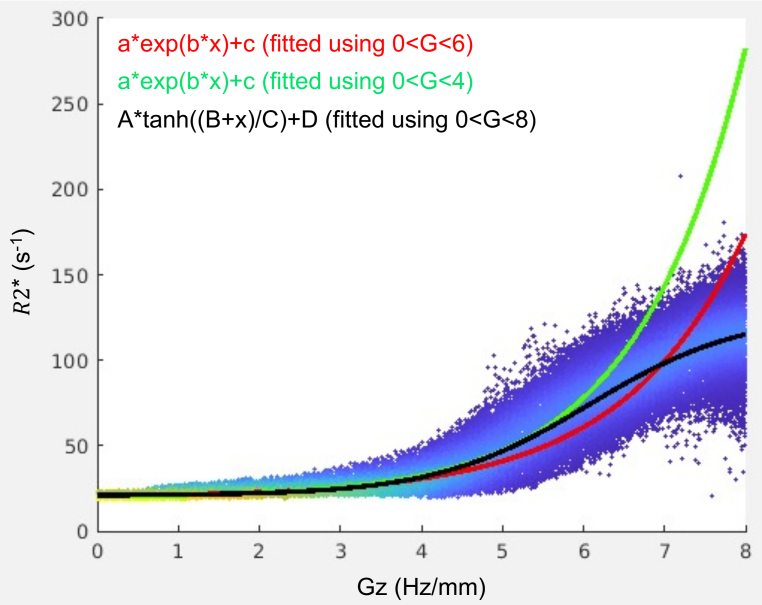


**Figure S14** Relationship between R2* and Gmag using voxel-wise data from 150 UK Biobank subjects, fitting with an exponential and tanh model. The data scatter displays median R2* measures averaged across Gmag bins, with the colour representing the scatter density (blue – low density, green – high density). As the majority of median Gmag estimates were in the range 0<Gmag<4 Hz/mm, the exponential model (*a*exp(b*x)+c)* was fit to data in the range of 0<G<4 Hz/mm (green line) and of 0<G<6 Hz/mm (red line). The tanh function (*A*tanh((B+x)/C)+D)* (black line) showed the best performance across the whole range, particularly when Gmag > 6 Hz/mm.

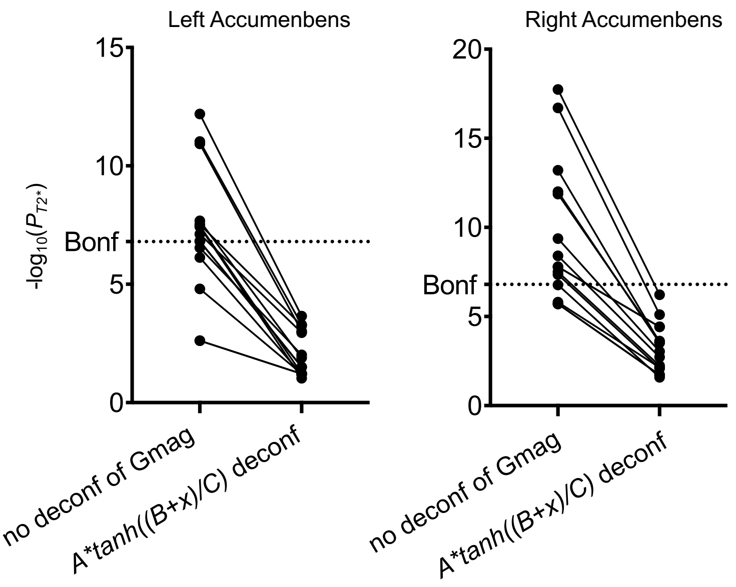


**Figure S15** Association $-\log_{10}P$ values between T2* accumbens IDPs and sinus/nasal-related phenotypes. Here, each point represents one association with one unique phenotype. These associations are considered spurious as accumbens regions are in the vicinity of the sinus and no previous literature has linked T2* in accumbens with sinus conditions. Deconfounding using "exp" or "tanh" model was able to dramatically reduce these associations to non-significant.

No association was found between the macroscopic field gradients and QSM data (**Fig. S16**), and thus this particular confound was only applied to T2* data.

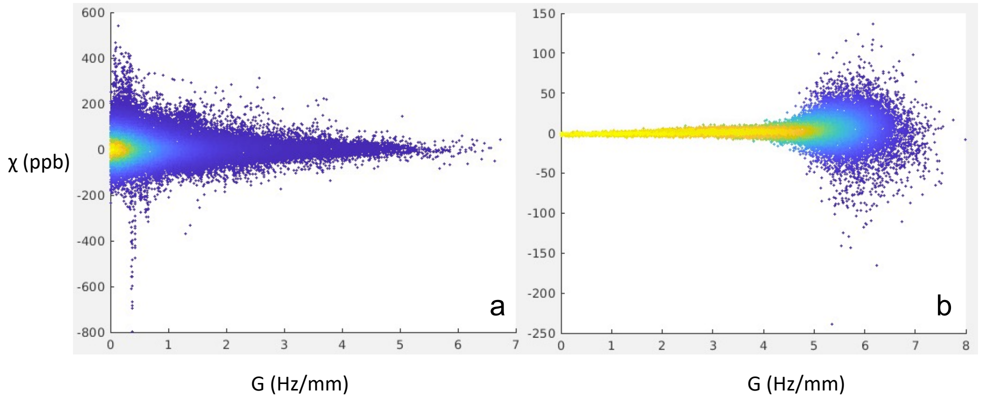


**Figure S16** Relationship between $\chi$ measures and macroscopic field gradient Gmag. **(a)** $\chi$ vs Gmag in voxel-wise data from 150 subjects and **(b)** median value in each bin of Gmag in **(a)**.

### Section 3: comparisons of phenotypic association between QSM and T2* IDPs

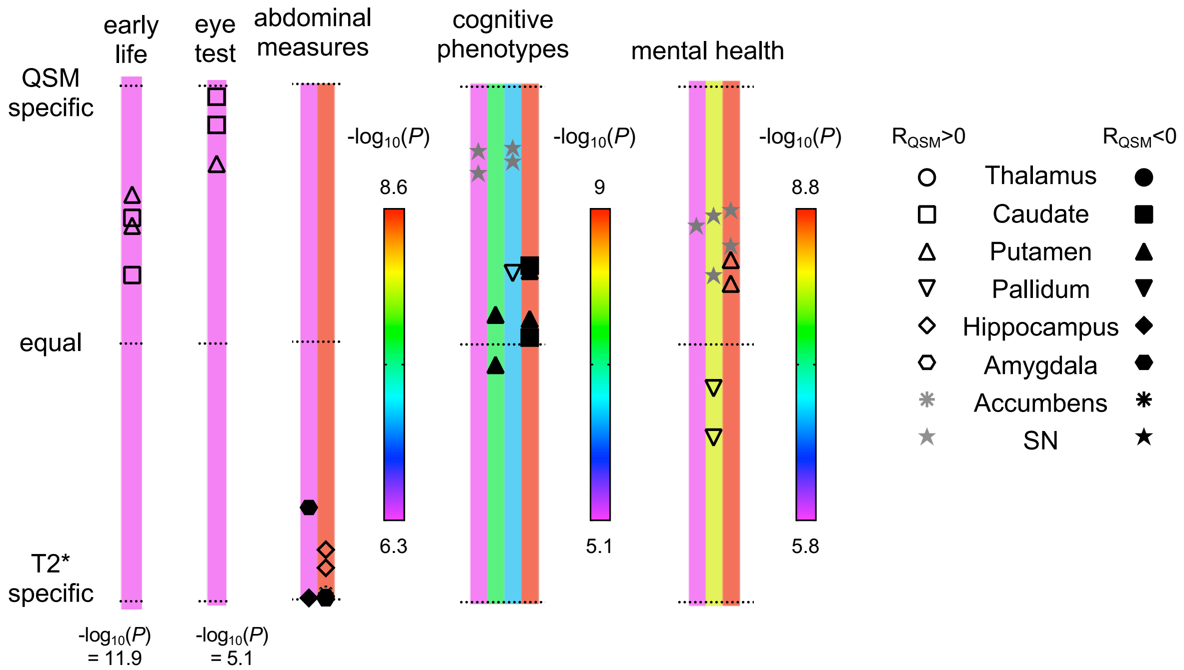


**Figure S17** Transformed Bland-Altman plot for the 5 categories that showed the smallest number of associations (early life factors, eye test, abdominal measures, cognitive phenotypes and mental health).

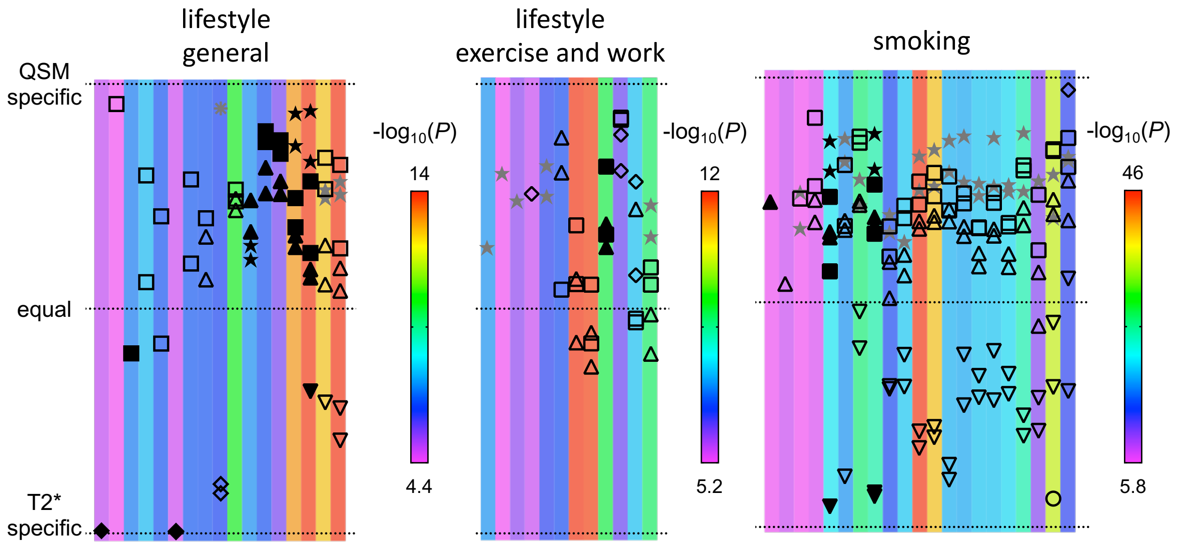


**Figure S18** Transformed Bland-Altman plot for lifestyle general, lifestyle exercise and work, smoking categories.

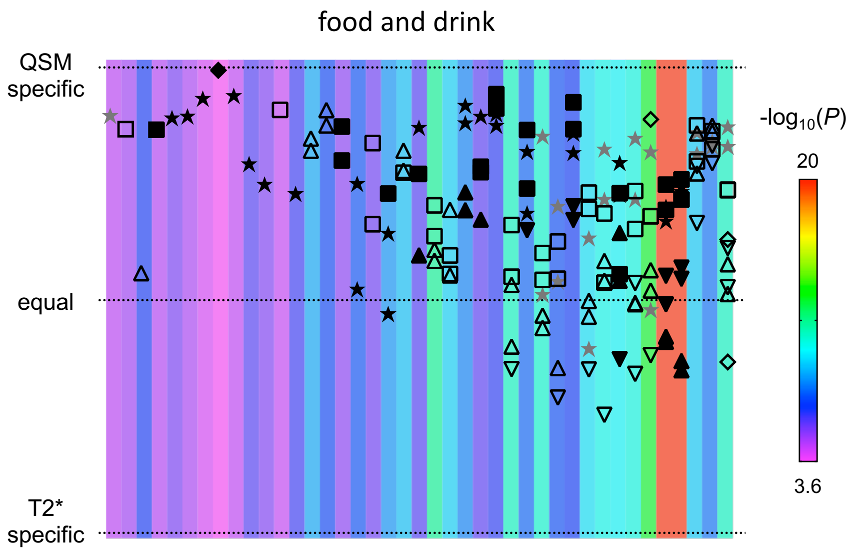


**Figure S19** Transformed Bland-Altman plot for the food and drink category.

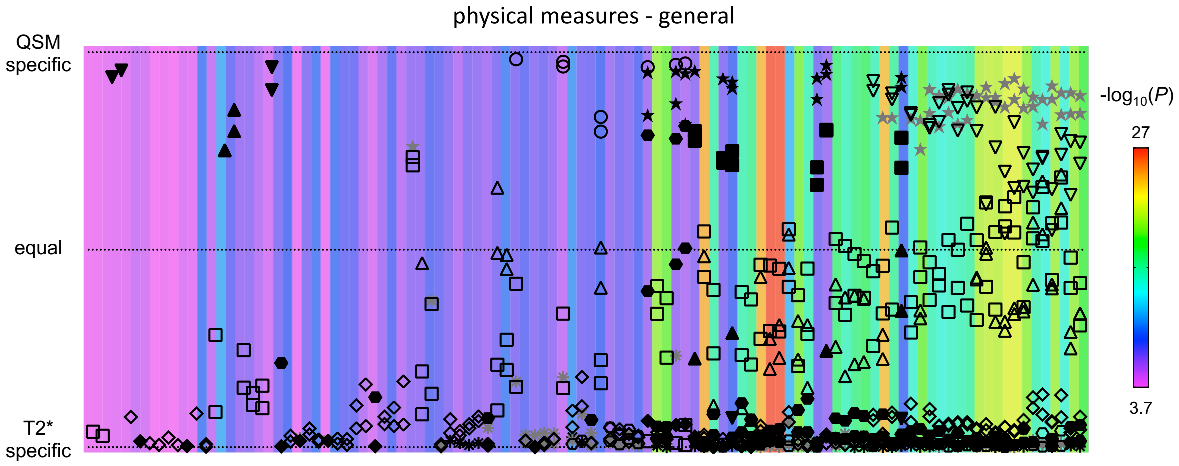


**Figure S20** Transformed Bland-Altman plot for the physical measures (general) category.

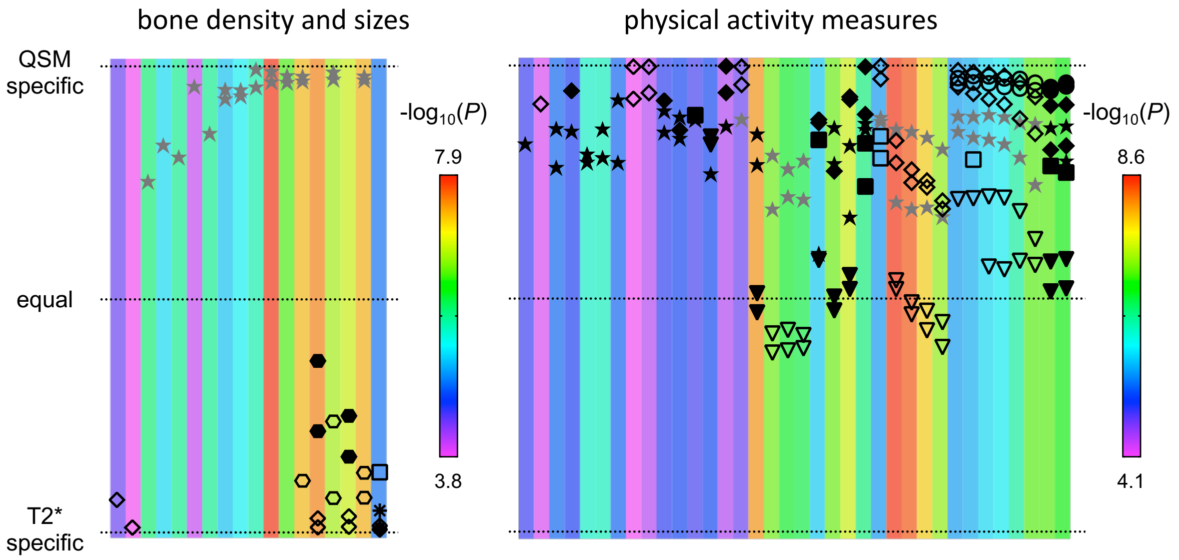


**Figure S21** Transformed Bland-Altman plot for bone density and physical activity categories.

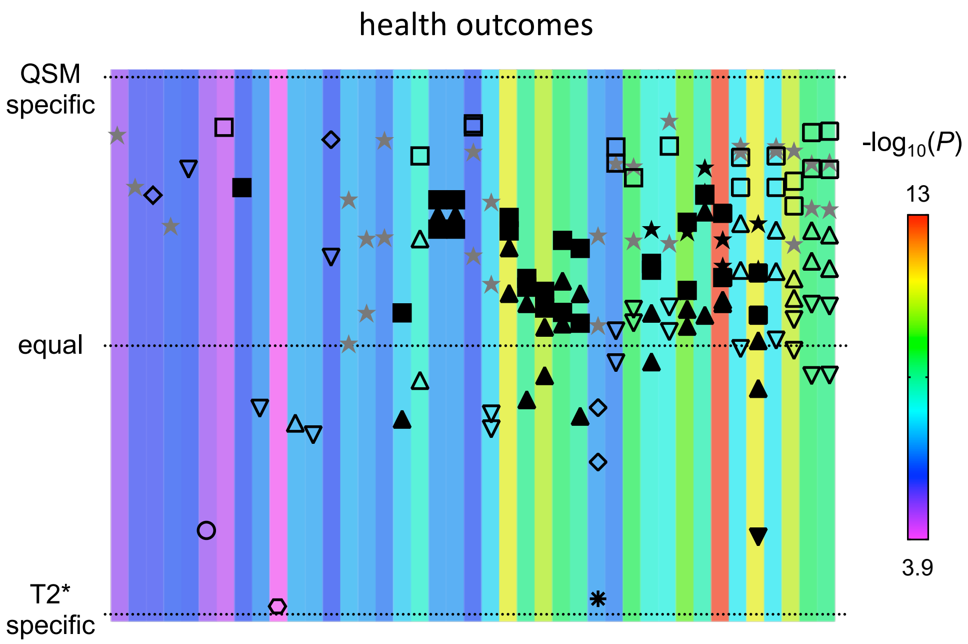


**Figure S22** Transformed Bland-Altman plot for the health outcomes category.

### Section 4: χ voxel-wise association maps for example associations in each phenotype category

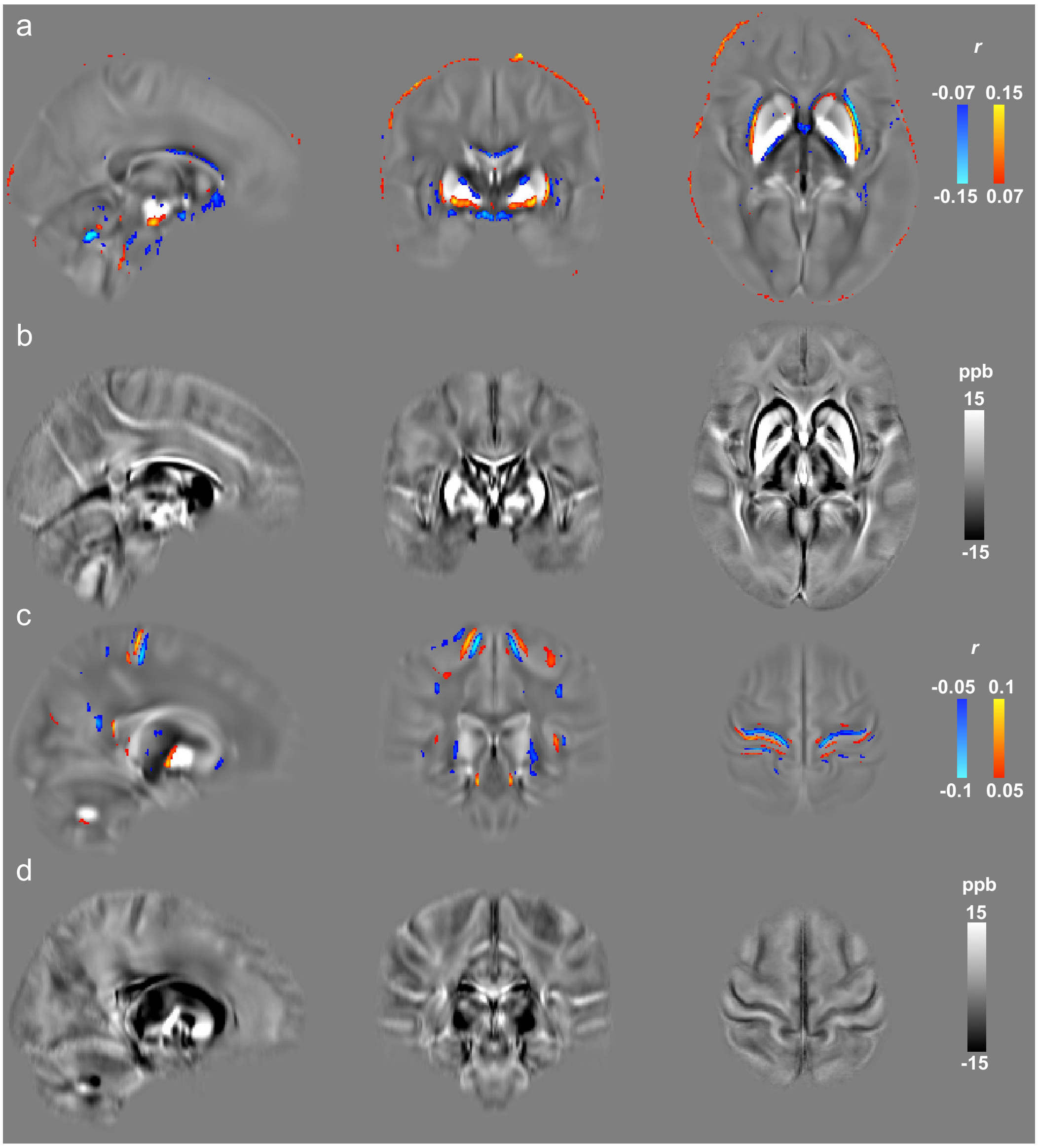


**Figure S23 (a)** Voxel-wise correlation map for *Total BMD*, with *r* values overlaid onto the susceptibility atlas. **(b)** Susceptibility aging atlas, generated by taking the difference between susceptibility maps from the youngest (< 52yo) and oldest (> 75yo) age groups in UK Biobank. Here, differences due to aging are largely driven by atrophy, displaying similar contrast to the regions of high *r* values in (a). **(c)** Voxel-wise correlations maps of *BMI* also demonstrate similar contrast to regions of brain atrophy, with **(d)** displaying the aging atlas in the same slices as (c).

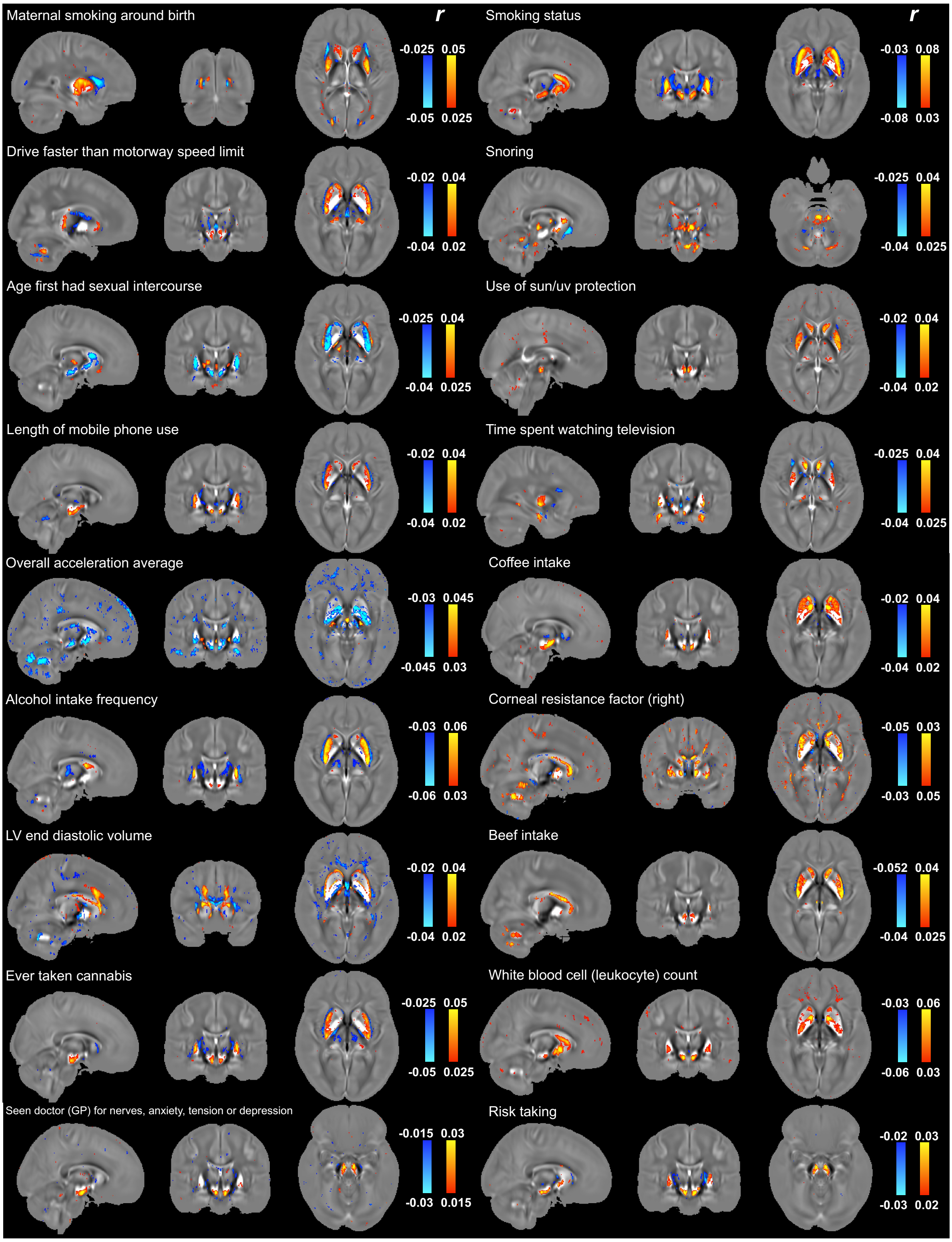


**Figure S24** Voxel-wise correlation maps with *r* values overlaid onto the susceptibility atlas.

### Section 5: Additional GWAS results for QSM and T2* IDPs

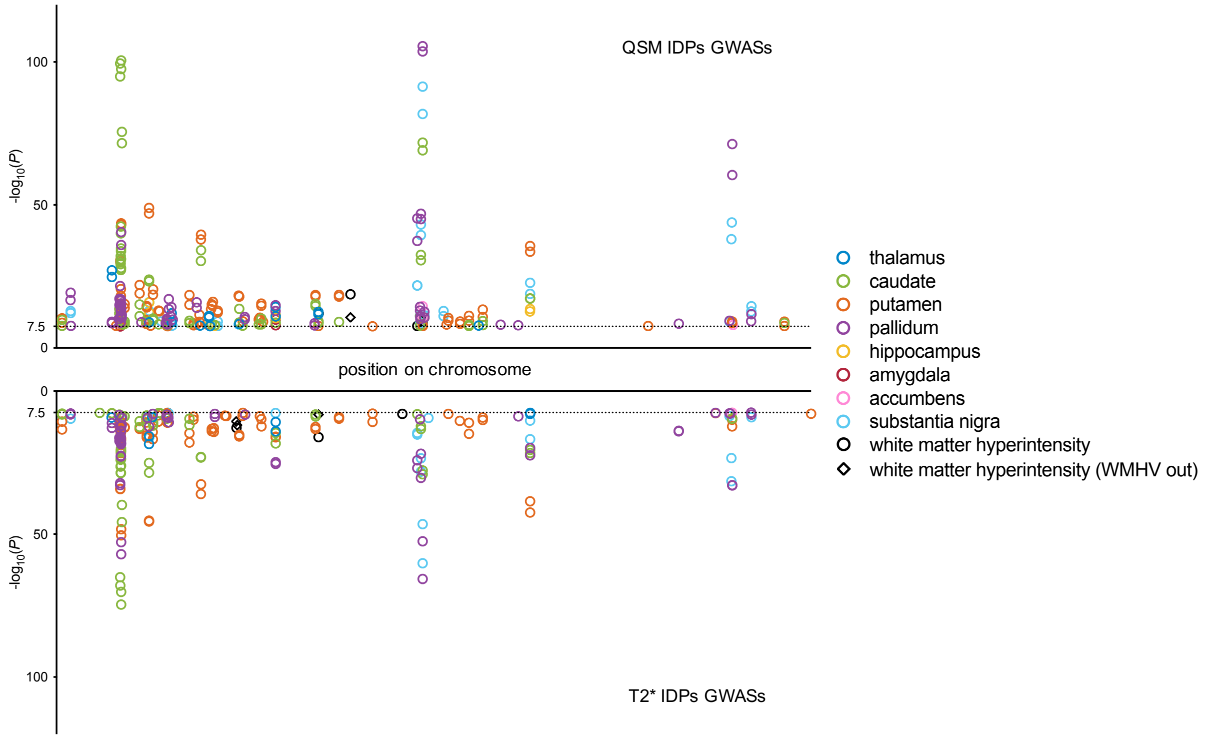


**Figure S25** Comparison of all lead genetic associations identified in GWASs of every QSM/T2* IDP. Each circle represents an association between a IDP and a lead genetic variant where x-axis showing variant's position on chromosome and y-axis gives its -$\log_{10}P$ in discovery cohort. QSM IDPs generally showed more genetic associations and higher -$\log_{10}P$ values compared to T2* IDPs.

Manhattan plots of every GWAS performed for QSM and T2* IDPs are shown below.

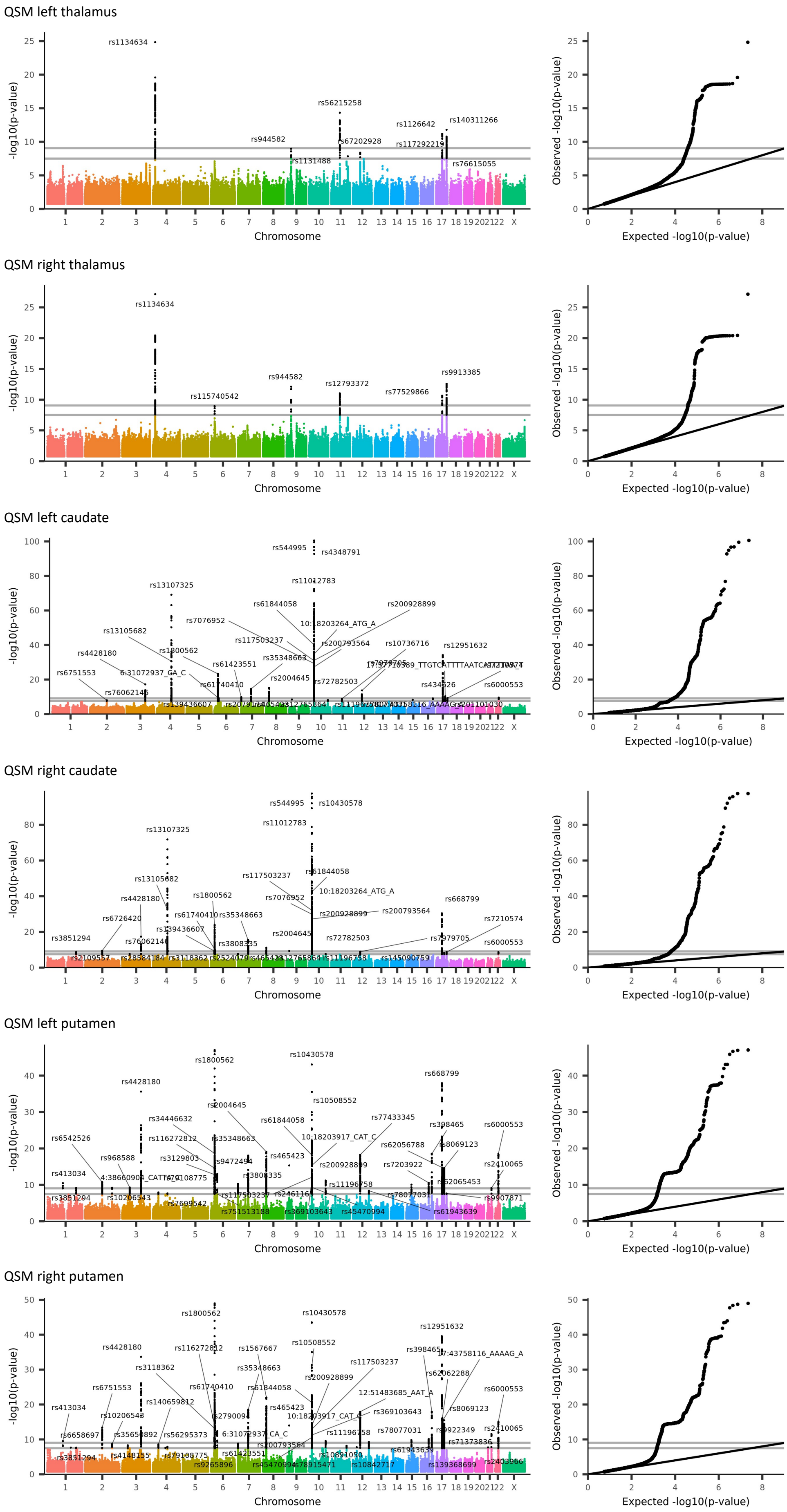
QSM left thalamus
rs1134634
rs56215258
rs944582
rs1126642
rs140311266
rs67202928
rs117292219
rs1131488
rs76615055
QSM right thalamus
rs1134634
rs944582
rs12793372
rs9913385
rs77529866
rs115740542
QSM left caudate
rs544995
rs4348791
rs11012783
rs13107325
rs61844058
rs200928899
rs7076952
10:18203264_ATG_A
rs13105682
rs117503237
rs200793564
rs10736716
rs12951632
rs4428180
rs1800562
rs35348663
rs61423551
rs6751553
6:31072937_GA_C
rs2004645
rs72782503
rs434326
rs6000553
rs61740410
rs76062146
rs139436607
QSM right caudate
rs544995
rs10430578
rs11012783
rs13107325
rs61844058
rs117503237
rs13105682
10:18203264_ATG_A
rs4428180
rs1800562
rs7076952
rs668799
rs200928899
rs200793564
rs6726420
rs61740410
rs35348663
rs7210574
rs139436607
rs2004645
rs3851294
rs76062146
rs72782503
rs7979705
rs6000553
rs3808335
rs2109557
rs145090759
QSM left putamen
rs10430578
rs1800562
rs668799
rs4428180
rs10508552
rs2004645
rs34446632
rs61844058
rs77433345
rs398465
rs6000553
rs6542526
rs116272812
rs35348663
10:18203917_CAT_C
rs62056788
rs8069123
rs968588
rs3129803
rs9472494
rs465423
rs200928899
rs7203922
rs2410065
rs413034
rs3808335
rs11196758
rs62065453
rs3851294
rs10206543
rs117503237
rs2461166
rs78077031
rs9907871
rs7699542
rs751513188
rs369103643
rs45470994
rs61943639
QSM right putamen
rs1800562
rs10430578
rs12951632
rs4428180
rs116272812
rs1567667
rs10508552
rs398465
17:43758116_AAAAG_A
rs3118362
rs35348663
rs117503237
rs200928899
rs62062288
rs6751553
rs61740410
rs61844058
12:51483685_AAT_A
rs6000553
rs413034
rs140659812
rs465423
rs8069123
rs10206543
rs2790094
10:18203917_CAT_C
rs369103643
rs6658697
rs35650892
rs56295373
6:31072937_CA_C
rs11196758
rs78077031
rs9922349
rs2410065
rs200793564
rs71373836
rs3851294
rs4148155
rs79108775
rs61423551
rs10891050
rs61943639
rs9265896
rs45470994
rs78915471
rs10842717
rs139368699
rs2403966
Chromosome
-log10(p-value)
Expected -log10(p-value)
Observed -log10(p-value)

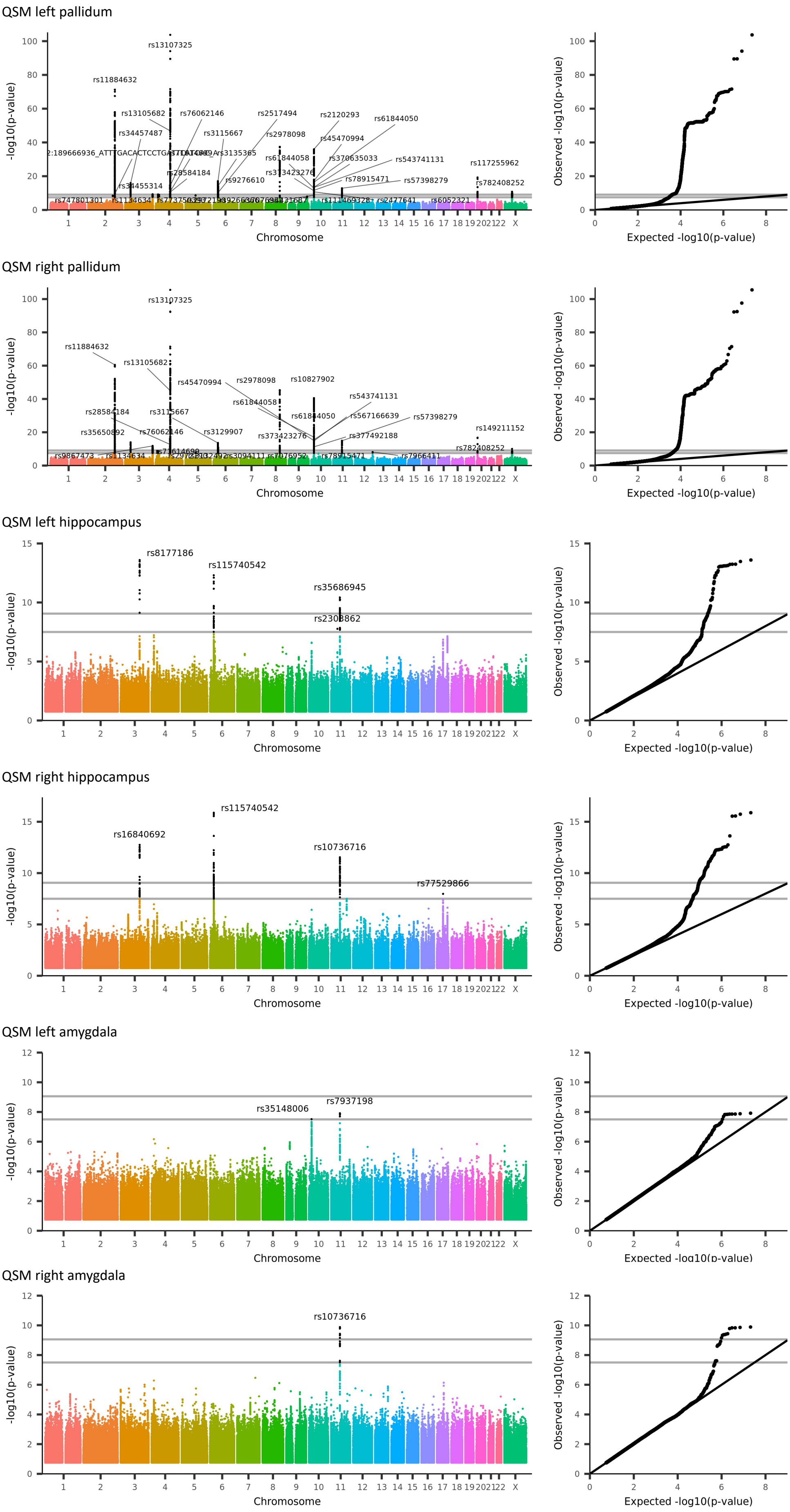

QSM left pallidum
rs13107325
rs11884632
rs13105682
rs76062146
rs2517494
rs2120293
rs61844050
rs34457487
rs3115667
rs2978098
rs45470994
rs3135365
rs61844058
rs370635033
rs543741131
rs117255962
rs28584184
rs373423276
rs9276610
rs78915471
rs57398279
rs782408252
rs34455314
rs747801301
rs1134634
rs2477641
rs6052321
-log10(p-value)
Chromosome
Observed -log10(p-value)
Expected -log10(p-value)
QSM right pallidum
rs13107325
rs11884632
rs13105682
rs45470994
rs2978098
rs10827902
rs543741131
rs61844058
rs28584184
rs3115667
rs61844050
rs567166639
rs57398279
rs35650892
rs76062146
rs3129907
rs373423276
rs377492188
rs149211152
rs782408252
rs9867473
rs1134634
rs7076952
rs78915471
rs7966411
-log10(p-value)
Chromosome
Observed -log10(p-value)
Expected -log10(p-value)
QSM left hippocampus
rs8177186
rs115740542
rs35686945
rs2308862
-log10(p-value)
Chromosome
Observed -log10(p-value)
Expected -log10(p-value)
QSM right hippocampus
rs115740542
rs16840692
rs10736716
rs77529866
-log10(p-value)
Chromosome
Observed -log10(p-value)
Expected -log10(p-value)
QSM left amygdala
rs7937198
rs35148006
-log10(p-value)
Chromosome
Observed -log10(p-value)
QSM right amygdala
rs10736716
-log10(p-value)
Chromosome
Observed -log10(p-value)
Expected -log10(p-value)

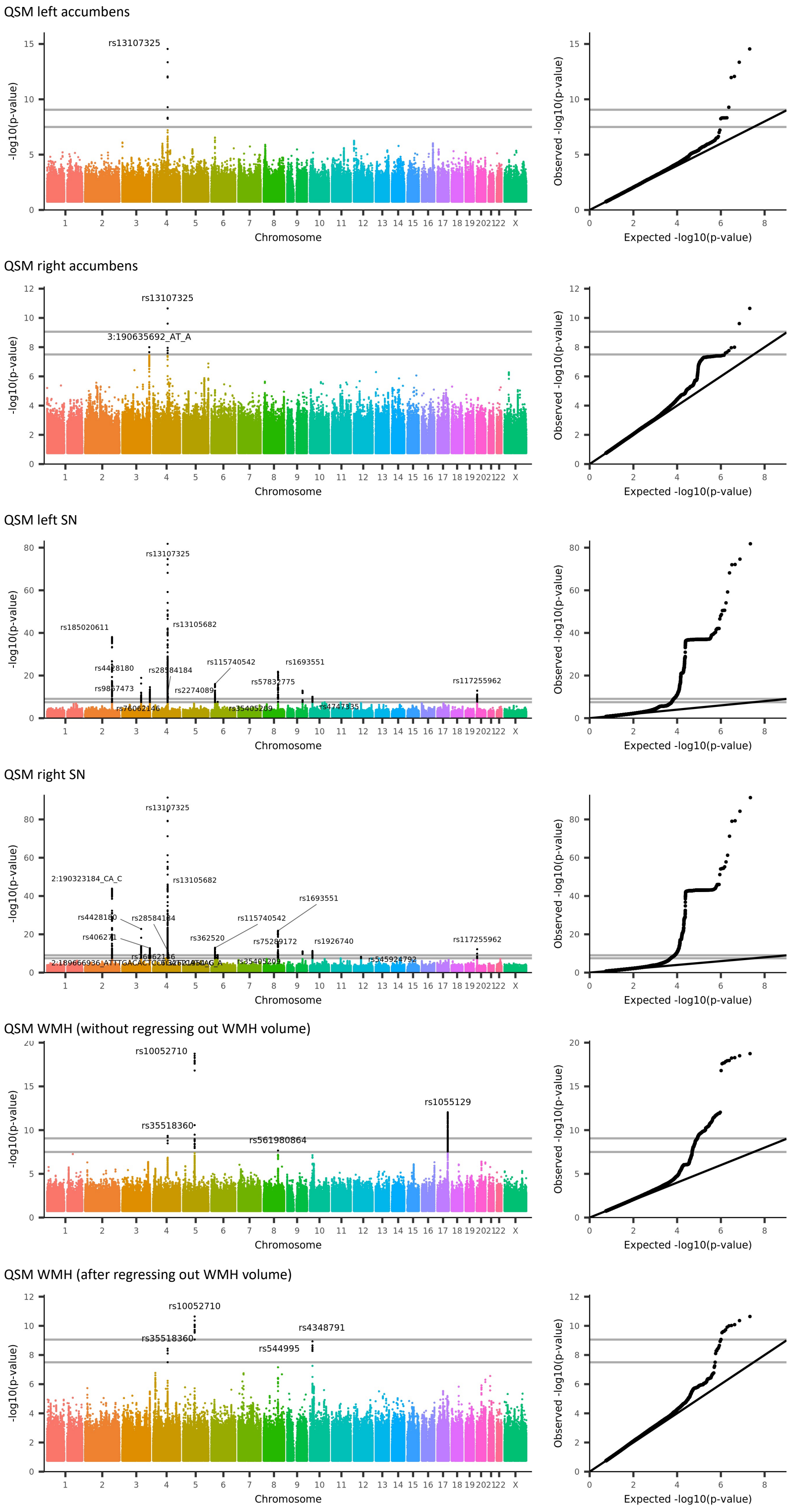
QSM left accumbens
rs13107325
-log10(p-value)
Chromosome
Observed -log10(p-value)
Expected -log10(p-value)
QSM right accumbens
rs13107325
3:190635692_AT_A
QSM left SN
rs13107325
rs13105682
rs185020611
rs4428180
rs28584184
rs115740542
rs1693551
rs57832775
rs117255962
rs2274089
rs35405209
rs4747335
QSM right SN
rs13107325
rs13105682
2:190323184_CA_C
rs1693551
rs4428180
rs28584184
rs115740542
rs362520
rs1926740
rs117255962
rs35405209
rs545924792
QSM WMH (without regressing out WMH volume)
rs10052710
rs1055129
rs35518360
rs561980864
QSM WMH (after regressing out WMH volume)
rs10052710
rs4348791
rs35518360
rs544995

T2* left thalamus

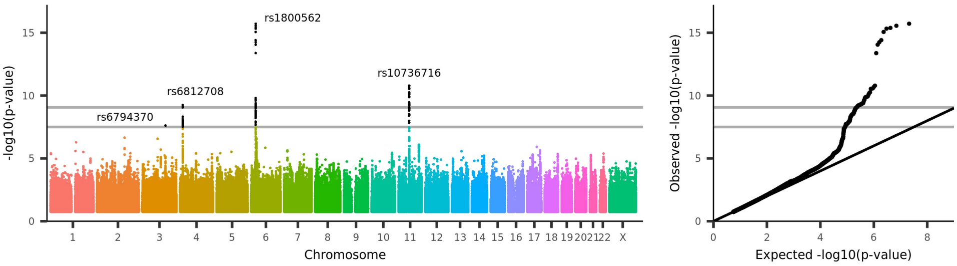


T2* right thalamus

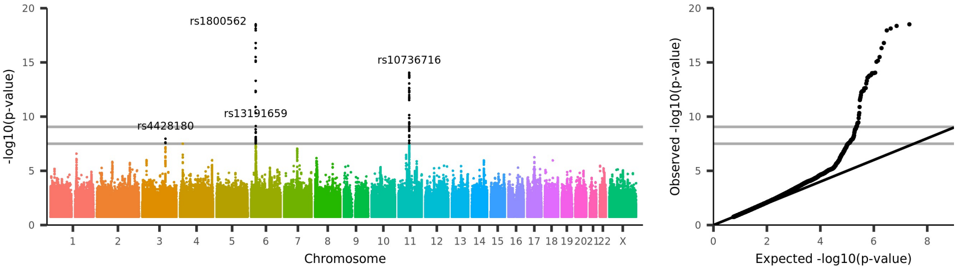


T2* left caudate

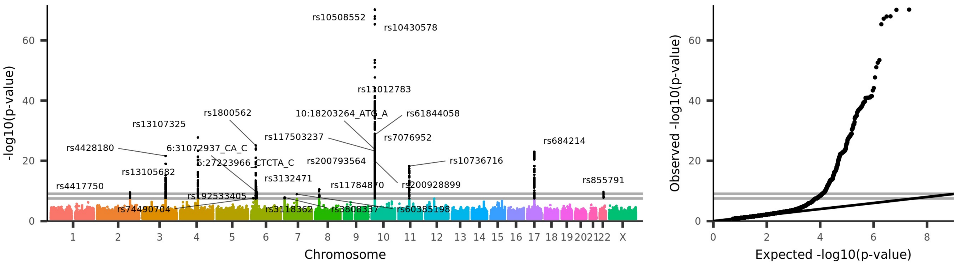


T2* right caudate

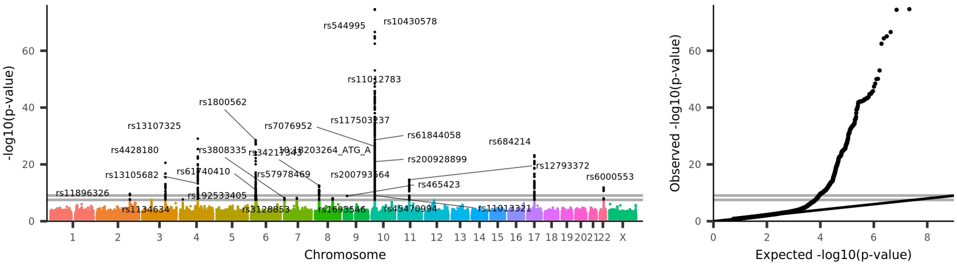


T2* left putamen

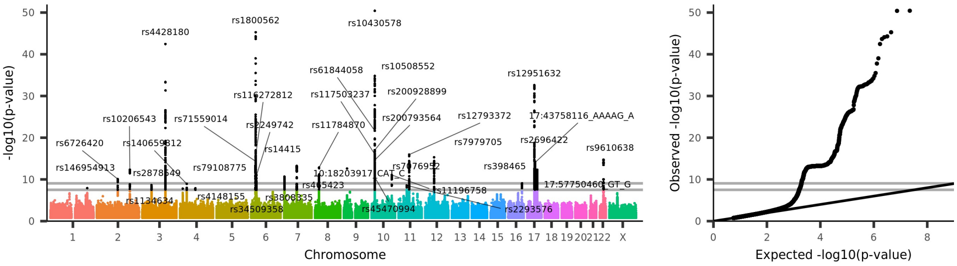


T2* right putamen

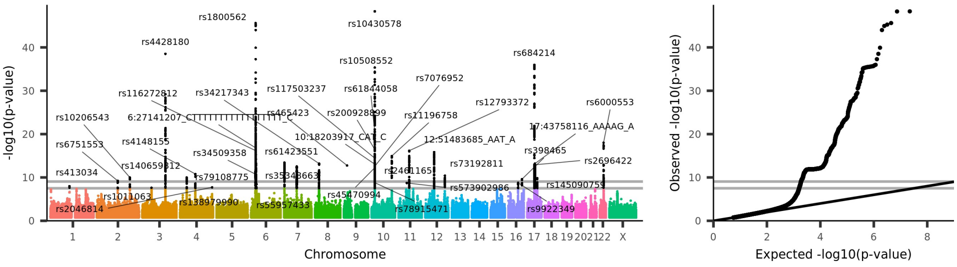

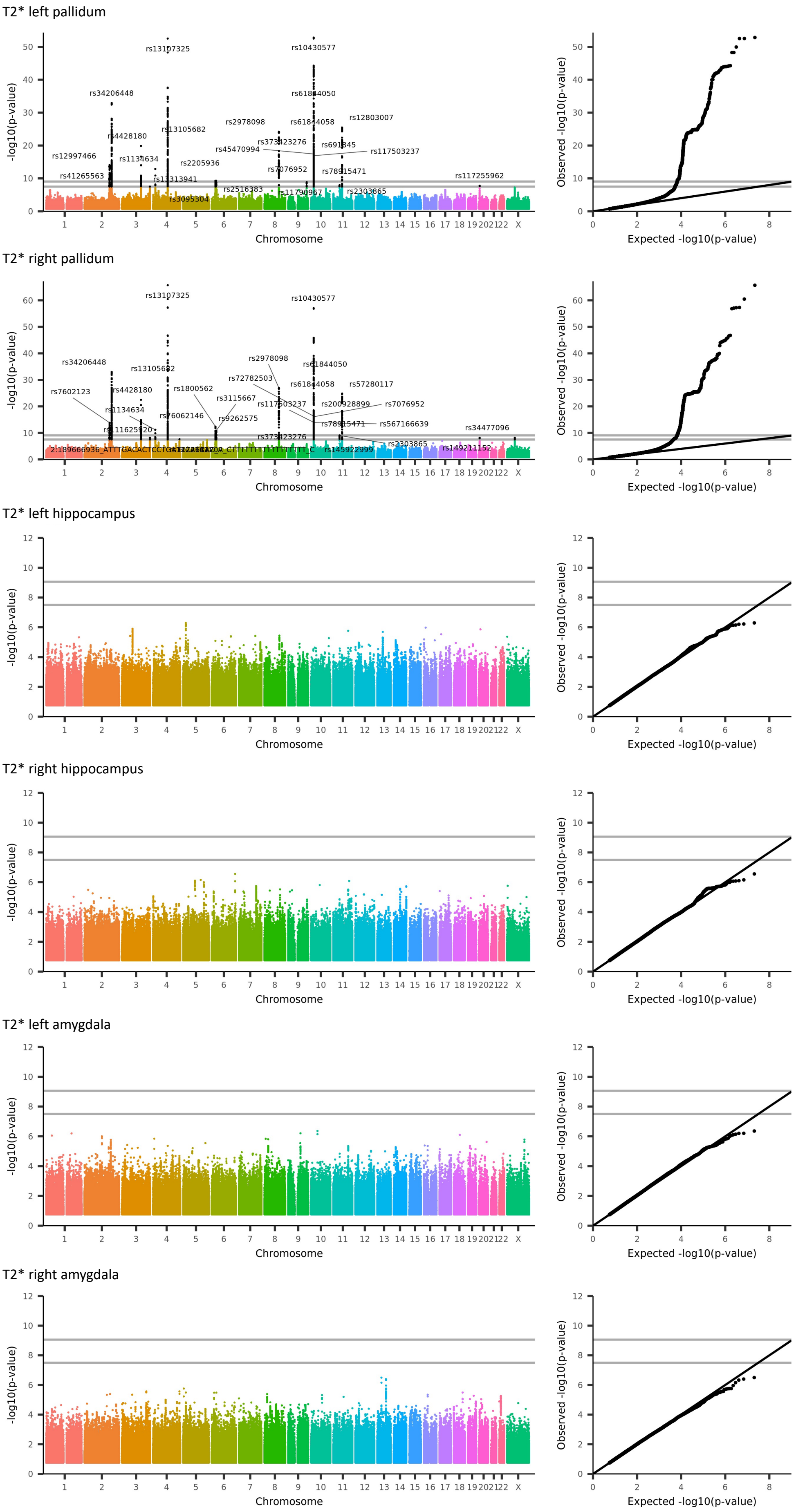
T2* left pallidum
rs13107325
rs10430577
rs34206448
rs61844050
rs12803007
rs13105682
rs2978098
rs61844058
rs4428180
rs373423276
rs691345
rs45470994
rs117503237
rs12997466
rs1134634
rs2205936
rs7076952
rs78915471
rs41265563
rs13313941
rs117255962
rs2516383
rs11790967
rs2303865
rs3095304
T2* right pallidum
rs13107325
rs10430577
rs2978098
rs34206448
rs13105682
rs61844050
rs72782503
rs57280117
rs7602123
rs4428180
rs1800562
rs61844058
rs3115667
rs117503237
rs200928899
rs7076952
rs1134634
rs76062146
rs9262575
rs78915471
rs567166639
rs111625920
rs34477096
rs373423276
rs2303865
rs145922999
rs149211152
T2* left hippocampus
T2* right hippocampus
T2* left amygdala
T2* right amygdala
-log10(p-value)
Chromosome
Observed -log10(p-value)
Expected -log10(p-value)

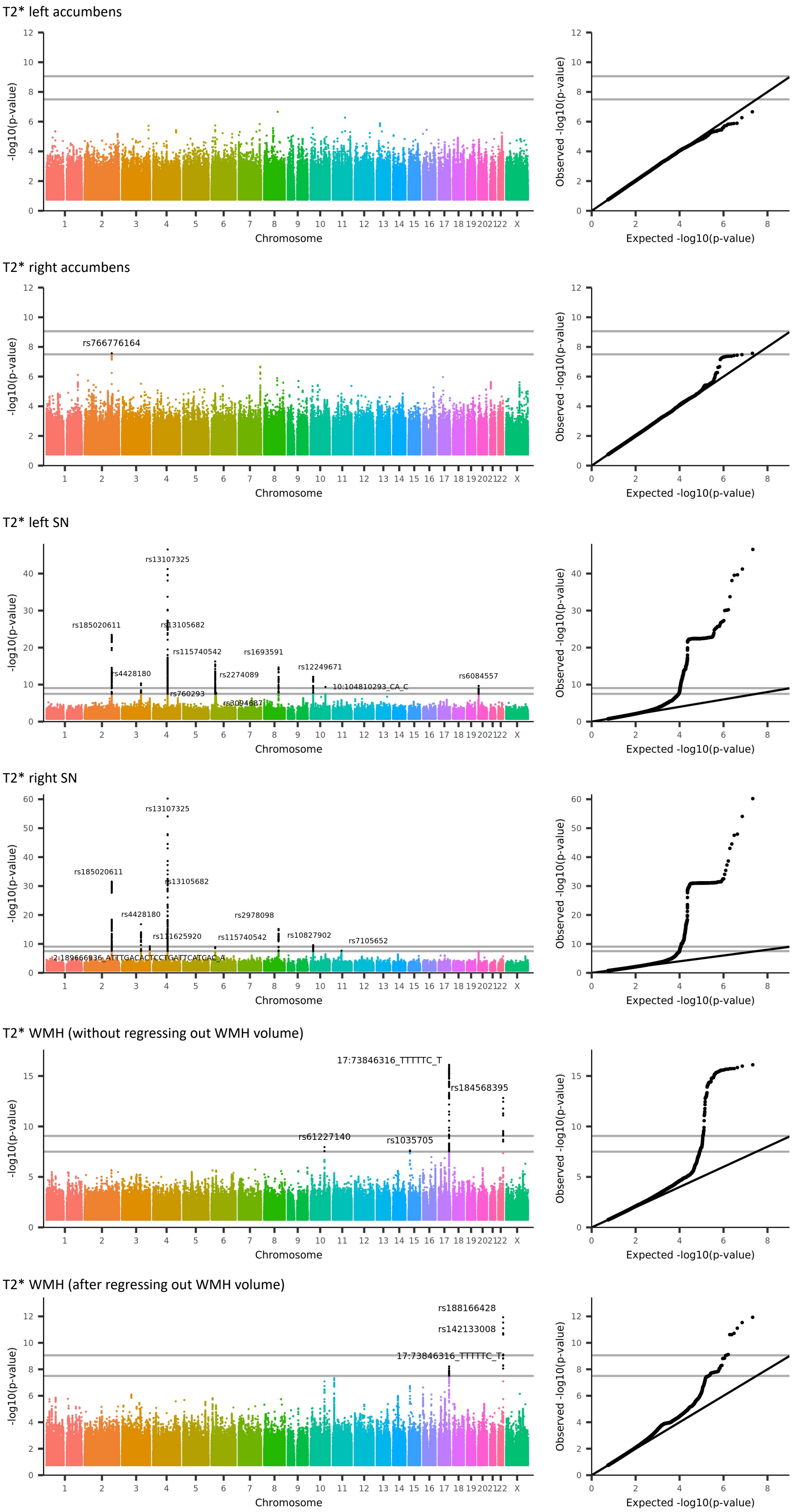

T2* left accumbens
-log10(p-value)
Chromosome
Observed -log10(p-value)
Expected -log10(p-value)
T2* right accumbens
rs766776164
T2* left SN
rs13107325
rs185020611
rs13105682
rs115740542
rs1693591
rs12249671
rs4428180
rs2274089
10:104810293_CA_C
rs6084557
rs760293
T2* right SN
rs13107325
rs185020611
rs13105682
rs4428180
rs2978098
rs111625920
rs115740542
rs10827902
rs7105652
T2* WMH (without regressing out WMH volume)
17:73846316_TTTTTC_T
rs184568395
rs61227140
rs1035705
T2* WMH (after regressing out WMH volume)
rs188166428
rs142133008
17:73846316_TTTTTC_T

### Section 6: Associations between c or T2* and genetic variants not directly related to myelin, iron and calcium homeostasis

Associations between QSM and variants in genes related to myelin, iron and calcium were expected, since they are all known to affect brain c[19]. However, many of the associations identified in this study could not be directly related to none of these pathways. A few notable examples were associations with genes encoding extracellular matrix proteins, transcription factors and proteins related to immune response.

We observed associates with two genes related to the extracellular matrix, COL3A1 and VCAN. Both QSM and T2* in the globus pallidum and substantia nigra were associated with variants related to the COL3A1 gene (cluster 9, peak variant 2:189666936_ATTTGACACTCCTGATTCATCAC_A, P=3.07*E-10). COL3A1 encodes the type III collagen, a fibrillar-forming collagen that is a major component of the extracellular matrix in a variety of organs in adults. It's also expressed throughout embryogenesis, and is considered to play a central role in cerebral cortex development. Patients with mutations in COL3A1 have a variety of connective tissue anomalies and can also present profound brain anomalies in both grey and white matter[20].

T2* in the globus palidus and QSM in the hippocampus were also associated with variants in the CD82 gene, including a synonymous exonic variant (cluster 62, rs2303865, P=9.42*E-10). CD82 encodes a membrane glycoprotein highly expressed in myelinating oligodendrocytes with multiple roles, such as metastasis suppression, immune response, and in the development of oligodendrocytes and maturation of oligodendrocyte precursors. It may regulate myelin proteins gene transcription or stabilize protein levels[21,22]. This same variant has been previously associated to white matter microstructure measurements [23]. Interestingly, in the projected maps, both GFAP and CD82 were associated with QSM in most of the white matter tracts.

Surprisingly, T2* and QSM IDPs were associated to many genetic variants related to transcription factors. Transcription factors regulate the transcription of DNA to RNA and can modulate the expression of multiple genes. It is challenging to anticipate how variants in transcription factors would lead to changes in T2* or QSM in the brain. As an example, QSM in the putamen and in substantia nigra were associated with a variant in the RUNX2 gene (cluster 33, rs9472494, P=1.02*E-13). This gene encodes a transcription factor that plays a major role in osteoblastic differentiation and skeletal morphogenesis, but that is expressed in many tissues including the brain[24]. In the nervous system, RUNX2 acts on its development, regeneration and repair, regulating multiple mechanisms in neurons and glia[25,26], but it can also promote ectopic vascular biomineralization[27]. Down-regulation of RUNX2 signalling has been reported in the dorsolateral prefrontal cortex of patients with schizophrenia[28], as well as reduced expression of RUNX2 in the hippocampus of patients with bipolar disorder[29]. In our study, the association between QSM and variants in RUNX2 could be a consequence of alterations in multiple pathways, such as an increase in vascular calcifications or differences in brain development leading to changes in tissue microstructure.

Finally, a high number of associations, including associations with very low p-values, were related to the gene MRC1 (or MRC1L1), with seven clusters and 78 associations being related to this gene (T2* and QSM in the globus pallidum, caudate, putamen, in addition to QSM in the amygdala and WMH; strongest associations in cluster 48, rs544995, P=3.18*E-100). MRC1 encodes a molecular scavenger protein, also known as CD206, that mediates the endocytosis of glycoproteins and is primarily related to immune response. It acts by clearing harmful glycoconjugates, enzymes, hormones, cell membranes, extracellular matrix components, and micro-organisms through recognition of their carbohydrate structures[30,31]. It mediates many roles, including clearance of inflammatory molecules[31], remodelling of the extracellular matrix 52, immune response and cavity and scar formation after injury in the CNS[32], and is known to be expressed by astrocytes and microglia in the brain[33]. In autism spectrum disorder, MRC1 is overexpressed in the white matter[34], and the variants rs544995 has been related to sarcoidosis, a chronic inflammatory disease[35].

How variants in these genes could affect tissue magnetic susceptibility and T2* is not clear. Indirect effects via regulation of iron or calcium homeostasis, brain development, plasticity and myelination, immune response, regulation of cell cycle, and scavenging of harmful molecules, could result in the accumulation of iron and/or calcium or in differences in the brain microstructure, for example.

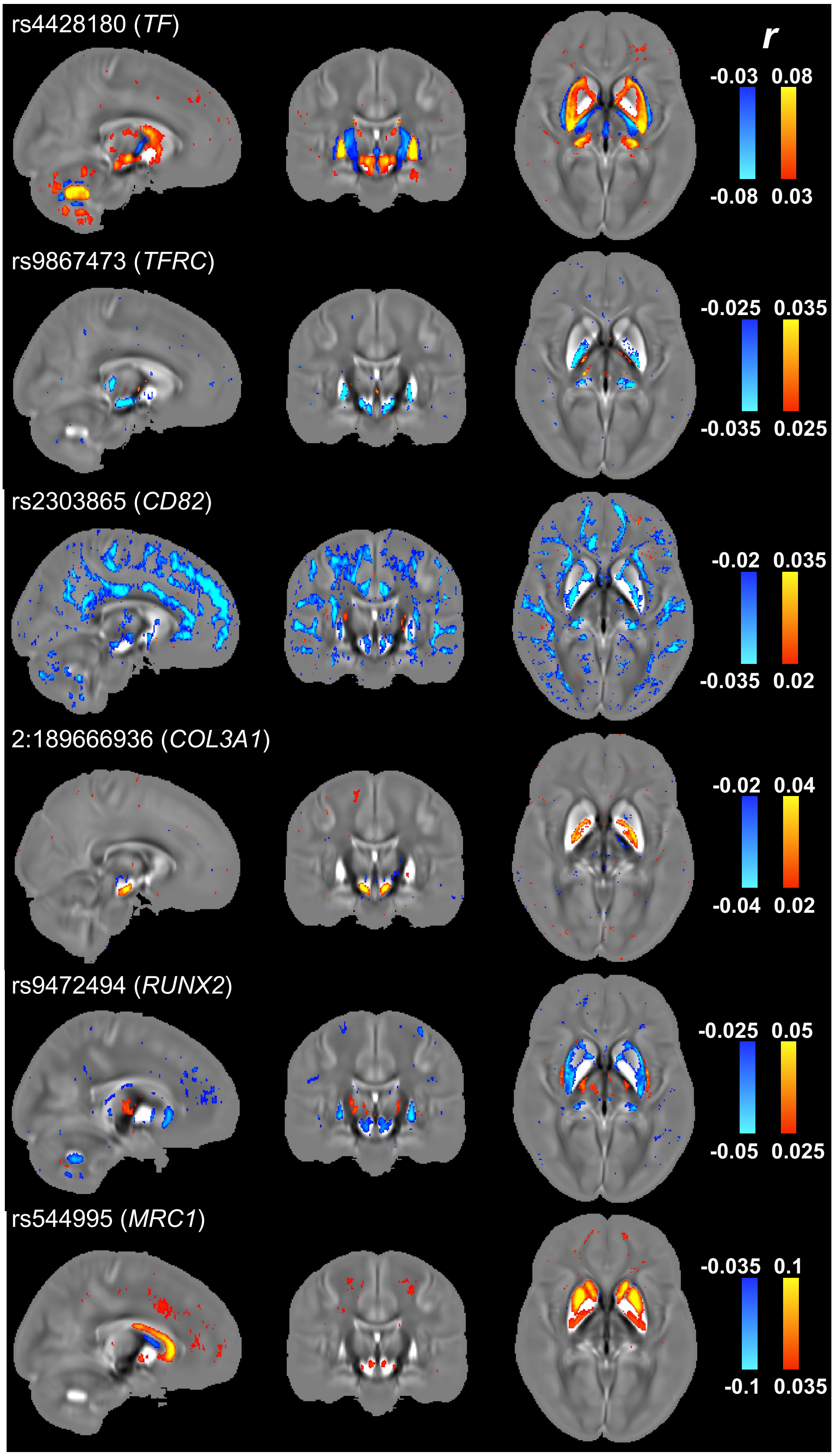


**Figure S26** Voxel-wise correlation maps of 6 top genetic variants, with *r* values overlaid onto the susceptibility atlas.
